# Failure to learn during roving, analysing the unsupervised bias hypothesis

**DOI:** 10.1101/383398

**Authors:** David Higgins, Michael Herzog

## Abstract

We examine the unsupervised bias hypothesis [11] as an explanation for failure to learn two bisection tasks, when task sequencing is randomly alternating (roving). This hypothesis is based on the idea that a covariance based synaptic plasticity rule, which is modulated by a reward signal, can be biased when reward is averaged across multiple tasks of differing difficulties. We find that, in our hands, the hypothesis in its original form can never explain roving. This drives us to develop an extended mathematical analysis, which demonstrates not one but two forms of unsupervised bias. One form interacts with overlapping task representations and the other does not. We find that overlapping task representations are much more susceptible to unsupervised biases than non-overlapping representations. Biases from non-overlapping representations are more likely to stabilise learning. But this in turn is incompatible with the experimental understanding of perceptual learning and task representation, in bisection tasks. Finally, we turn to alternative network encodings and find that they also are unlikely to explain failure to learn during task roving as a result of unsupervised biases. As a solution, we present a single critic hypothesis, which is consistent with recent literature and could explain roving by a, much simpler, certainty normalised reward signalling mechanism.

## 1 Introduction

Performance in many perceptual tasks improves with practice, a paradigm strongly associated with learning. Some well known visual perceptual learning tasks include orientation discrimination [19], contrast discrimination [33], detection of moving dot direction [3, 16], vernier offset discrimination [20] and bisection offset discrimination [4]. In each case, repeated training leads to an improvement in subjects’ threshold of discrimination on the task. For a review, see for example [7].

Bisection offset discrimination involves the presentation of three vertical lines, with a fixed distance separating the two outer lines (see **Figure 1,a**). Subjects must determine whether the central line is offset to the ‘left’ or to the ‘right’ of the true (not visible) bisector of the space, in a two alternative forced choice scenario. Given performance feedback, human subjects can decrease their threshold of discrimination over time which leads to better performance. Increasing the outer line distance by ≥50% leads to performance on each which suggests no transfer, or occlusion, in learning, takes place, from one outer line distance to the other [22]. This has led to these differing outer line separations being referred to as different ‘tasks’ in the literature.

**Figure 1:**
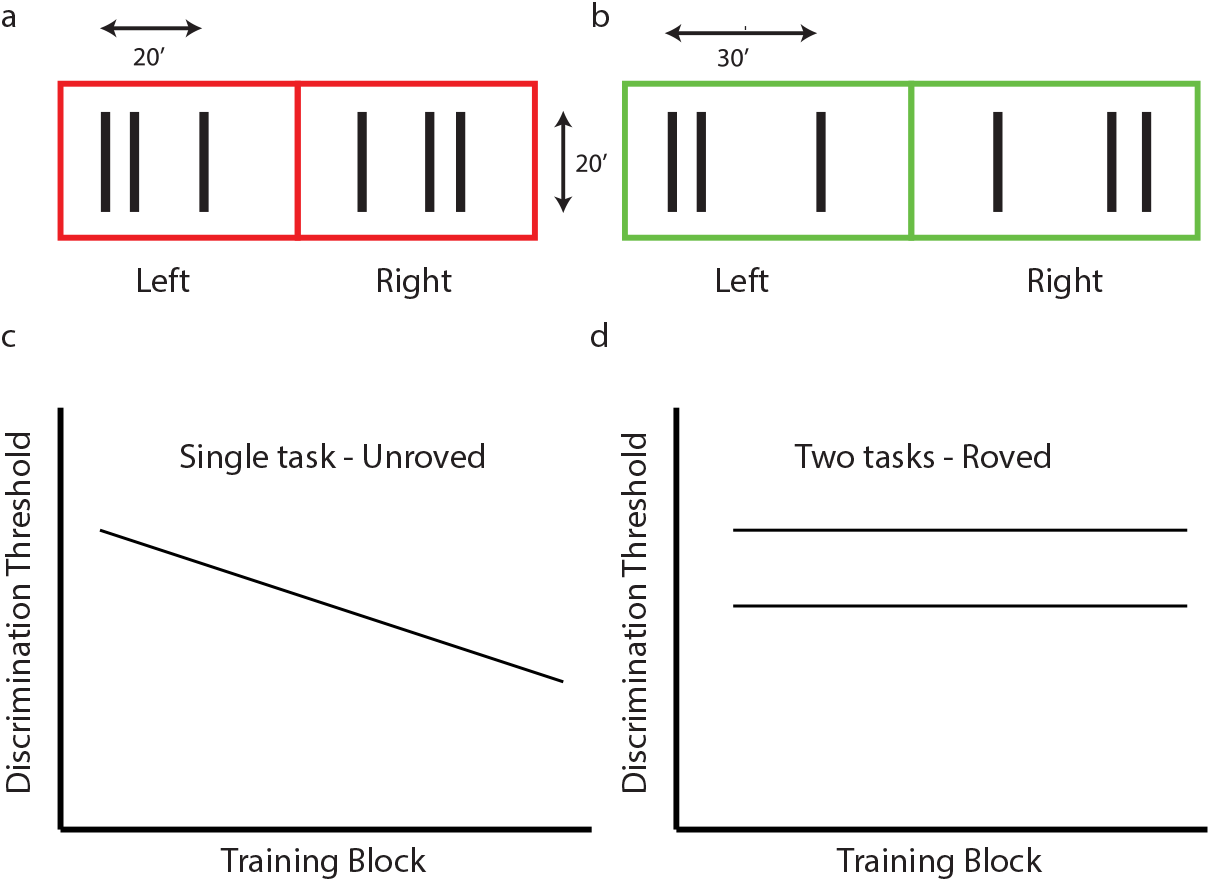
Sketch of bisection learning paradigm and results. (a and b) Tasks are identified by their outer line separation (20’ or 30’), highlighted here in red or green. Three vertical bars are presented and subjects must determine whether the central bar is to the left or the right of the true bisector of the space between the outer bars. (c) With feedback, subjects can improve their discrimination threshold on either task over time. (d) However, randomly switching between tasks on a trial-to-trial basis (roving) prevents improvements in the discrimination threshold from occurring.

Roving refers to the random sequencing of tasks in a perceptual learning experiment [16, 22]. In the case of bisection tasks this means that on a trial-to-trial basis, either the ‘narrow’ or the ‘wide’ outer line separation task is presented (see **Figure 1, a and b**). Within the given task the selection of a left or right stimulus, and its offset, is determined by the individual task paradigm. This means that stimuli are typically equiprobable (left-right) and a fixed offset degree, based on subjects’ initially measured thresholds of discrimination.

Roving of bisection offset discrimination tasks, with sufficiently large differences in their outer line separation, has been shown to impede learning [22] (compare **Figures 1,c and d**). In [22] the outer lines were not always presented in a fixed position, per task, but rather moved laterally a random degree within a given bounding area. This randomised the location of the discrimination stimulus on the retina per task. Using a fixed stimulus location, it was later shown [29] that the moving of the stimulus location was not necessary for the impediment to learning. Rather roving of the two tasks alone was responsible for the failure to learn.

Parkosadze et al. (2008) [23] pushed the roving-bisection learning paradigm to new limits, extending the number of repetitions from approximately 1,100 trials [22] to 18,000 trials. They showed that, following a significant period of worsening performance (at least 3,600 trials) the subjects’ performance eventually went through a transition and learned, rather quickly, both tasks, to levels typically seen when tasks are learned separately.

A number of approaches have used to model perceptual learning. Signal detection theory has been applied to distinguish between learning which is occurring at the perceptual level from that at the decision criterion level [1]. Channel noise theories have allowed the systematic experimentation with noise to constrain the possible sources of noise in a circuit for perceptual learning [6]. Artificial neural network models have been used to isolate the premises of perceptual learning and check their logical consistency [14]. Meanwhile, models of the early visual processing system have attempted to draw links from biologically inspired models to results of perceptual learning experiments [34, 27]. A review of modelling approaches can be found in [31].

In 2012, Herzog et al. [11] presented a theoretical explanation, including a model, of the failure to learn during roving based on the concept of an *unsupervised bias* in a reward-modulated synaptic plasticity rule. The unsupervised bias is a concept elaborated by Frémaux et al. (2010) [8] involving the *over-* or *under*estimation of reward in a reward-modulated covariance-based synaptic learning rule. In such a scenario, Frémaux et al. showed that correctly estimating the expectation of the reward in a nominally Temporal Difference (TD) learning scenario leads to much more stable learning outcomes. The *mis*-estimation of the correct expectation of reward leads to a potential systematic bias in the weight update rule, which in turn leads to worse task performance when compared with a reward learning rule which does not suffer from any such bias. A deeper explanation of this concept is provided in **Section 2.1.1**.

The theory of roving presented in [11] derives an unsupervised bias in the learning rule from the presence of a single reward predictor in the system. This reward prediction (typically referred to as *critic* in the reinforcement learning literature [28]) systematically over-estimates performance on one of the bisection tasks and under-estimates performance on the other. A feed-forward artificial neural network model was presented, in the paper, which successfully learns on a single bisection task and fails to learn when two tasks are roved.

In this paper, we examine the model from [11] in order to extend it to explain the transition to learning seen in [23]. In the process we find evidence which calls into question existing theories on how roving interferes with learning. We begin by examining the unsupervised bias hypothesis as it is presented in [11]. In **Section 2.1** we develop, and in **Section 3.1** we analyse, a simplified mathematical model of this hypothesis and find that it apparently cannot explain roving. In **Section 3.2** we extend the simplified analysis. In **Section 3.2.1** we show the mechanism by which the network presented in [11] actually fails to learn during roving. This mechanism turns out to be a form of unsupervised bias, but it cannot realistically explain roving. Numerical modelling results, presented in **Section 3.2.2**, highlight the existence of two forms of unsupervised bias only one of which directly interacts with the task representation. These results indicate that the unsupervised bias from a second (roved) task more typically aids in learning the first task. In **Section 4** we discuss alternative explanations for roving. In **Section 4.1** we examine whether any alternative task encodings can combine with the unsupervised bias hypothesis to explain roving and find that none are forthcoming. In **Section 4.2** we briefly discuss other approaches to modelling perceptual learning and whether they can be adapted to explain roving. Finally, in **Section 4.3** we present an alternative hypothesis, which takes the single critic theory from [11], combining it with recent experimental results, and uses it to provide a plausible explanation for the initial failure to learn during roving [22, 29] followed by the eventual transition to learning [23].

## 2. Methods

### 2.1. leSimplified model of the unsupervised bias hypothesis

In order to perform an analysis of the unsupervised bias hypothesis of failure to learn during task roving we need to build up our theory from a number of fundamental building blocks or assumptions. In **Section 2.1.1** we will clarify the meaning of the unsupervised bias hypothesis. In **Section 2.1.2**, we will define a reduced mathematical description of a network suitable for analysis of the unsupervised bias hypothesis. A basic prerequisite for the analysis to hold, is that we are in a reward prediction error learning framework [28]. This is justified with respect to existing modelling approaches for perceptual learning in **Section 4.2**.

The entire theory hangs on the idea of what the implications are of a single critic, which is incapable of distinguishing between two tasks. We will explore what this means for the Reward Prediction Error (RPE) required for learning. Our proof (presented in **Section 3.1**) of the insufficiency of this hypothesis hangs on this explanation and the generality of the representation assumptions.

#### 2.1.1. The unsupervised bias hypothesis

The unsupervised bias hypothesis is a theory of synaptic plasticity which has been derived from first principles [8]. Taking a covariance-based synaptic plasticity rule, it is possible to further add a reward-modulation term to the equations governing plasticity and obtain the following,

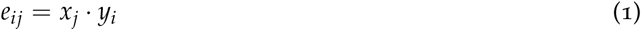

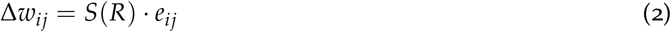

where *W_ij_* denotes the weight (synaptic efficacy) from presynaptic neuron *j* to postsynaptic neuron i. For the sake of comparison with Frémaux et al. [8], we separate the *elligbility trace, e_ij_*, into a separate variable. The structure is a single layered feedforward network, so notation is simplified with *x_j_* denoting presynaptic neuronal firing rates and *y_i_* postsynaptic neuronal firing rates. *S(R)* is a monotonic function of reward, derived from the global end-of-trial feedback signal, R. Note that, for comparison with roving experiments we are expressing everything in a trial-based formulation, however time continuous formulations are also available [8].

Now, assuming that the learning rate is sufficiently small, it is possible to take the expectation of the weight update rule, Δ*w*, with respect to both trials (time) and noise instances, and obtain the following,

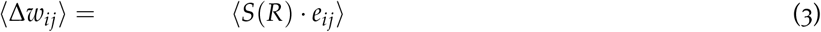

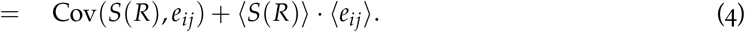

This result follows trivially from the definition of covariance. The covariance term, Cov(*S*(*R*),*e_ij_*), can be thought of as the pure learning term, or a *supervised* signal of learning. It corresponds perfectly to how the reward signal, in this case a Temporal Difference (TD) reward prediction error signal [28], coincides (covaries) with the network activity which led to the task performance and hence reward feedback signal. The two mathematical terms, 〈*S*(*R*)〉 and 〈*e_ij_*〉 combined, correspond to the unsupervised bias as examined by Frémaux et al. The word *unsupervised* refers to it being the part of the weight update rule which is left over after the covariance is mapped to the *supervised* signal. And *bias* refers to the clear effect of this set of mathematical terms on the dynamics of the synaptic weights, it is constant (across noise and trials) and, whether beneficial or negative in influence, does not correspond to the clearly desirable pure learning term.

**Figure 2:**
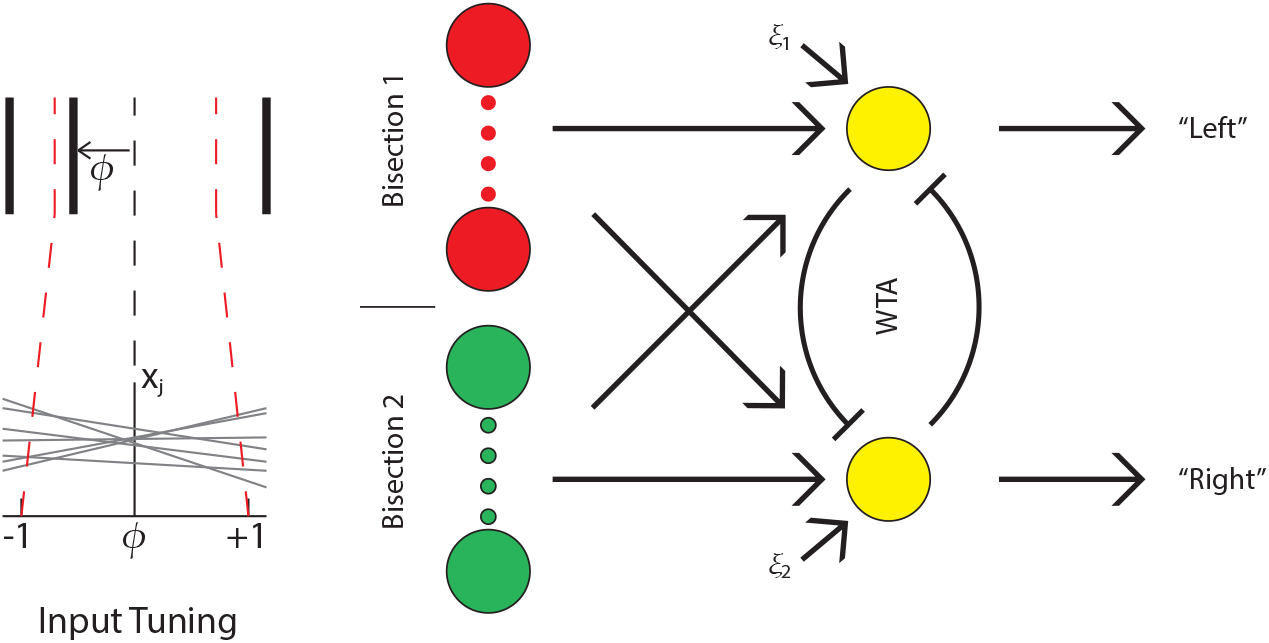
Herzog et al. (2012) network model to demonstrate the unsupervised bias hypothesis as an explanation for failure to learn during task roving. Different outer line distances (tasks) are passed via different pathways (red and green input neurons). Inputs are linearly tuned. All neurons are firing rate neurons. Input neurons are connected to output neurons via feedforward weights. Independent noise, ξ, is added to each of the output neurons. A winner-takes-all (WTA) mechanism on the output leads to a single output being active, per trial, which ‘chooses’ whether the input is left or right of the true bisection line.

#### 2.1.2. Mathematical description of the network

We can now develop a mathematical description of the network in order to analyse the suitability of the unsupervised bias hypothesis [11] as an explanation for failure to learn during roving.

Our first task is to clarify the input encoding. Herzog et al. used a linear encoding, *x*(*φ*) = *a* + *bφ*, where *a* ~ *N* (2,0.5) and *b* ~ *N* (0, *β*) per input neuron, with *β* regulating the difficulty of the classification task. For a presented stimulus, a true bisector is mapped to *φ* = 0, a left-of-centre middle line is mapped to*φ* ∊ [−1,0) and a right-of-centre middle line is mapped to *φ* ∊ (0,1]. This means that each input neuron encodes a single task presentation as approximately 2 + (*b* × distance from the true bisector), a linear encoding.

We have maintained this basic scheme, but have made some simplifications. These simplifications aid in the comprehension of the proof, but do not fundamentally change the computation being performed by the network. Firstly, we have completely removed all elements of randomness in the initialisation of our network. The firing rate for a true bisector is *a* = 0 for all input neurons. In terms of initial general computability of the underlying decision task, despite being biologically implausible, this poses no problems. This implementation also does not specifically conflict with the actual implementation of the network in [11]. We fix the slope of the tuning function to *b* = 1 for all neurons. This implies that a positive firing rate represents an input to the right (*φ* > 0) of the true bisector and a negative firing rate for all inputs to the left (*φ* < 0). This is massively simplifying the task representation but, again, is not computationally different from the network presented in [11].

We have considered whether more complicated input tuning functions are required than that presented in [11], or our simplification thereof. We find that while Gaussian or other smoothly changing functions certainly provide a degree of non-, or indeed super-linearity, which can aid in learning a classification (decision) they do not fundamentally change the classification to be performed. Furthermore, since we are mostly interested in the *failure* to learn during roving the linear representation can be considered a more interesting and challenging case. Previous attempts to explain bisection-task perceptual learning, have relied on physiologically known connectivity to lead to transfer/non-transfer of learning effects [34, 27]. In our case, we are testing only the effects of the unsupervised bias, allowing us to vastly simplify the encoding and thus the analysis. We will handle questions of alternative networks and encodings in **Section 4.1** and **4.2**.

Our other major change from [11] is that we now present only two input values to the system under analysis (*φ* ∊ { − 1,1}). This is actually closer to the protocols performed in the experiments and removes a small source of noise, where each input firing rate fluctuated according to the true distance of the input from the bisector. Our decision to use only the outer points on the interval, *φ* ∊ { − 1,1}, is motivated by a close examination of the effects on the Herzog et al. network of the use of intermediate values. In fact, these values are really only a source of noise on the input layer, which typically *smooths* the learning process. We are currently more interested in cases where the network *fails* to learn and thus are in no need of these intermediate values. Further, mathematically the use of these intermediate values does not change the classification problem being learned. Finally, as mentioned above, actual experimental procedure involves either presenting always the same angle left and right of the true bisector, as calibrated on a per subject basis, or within a tight range left and right, also on a per subject basis. In no case do experiments range the full distance from the true bisection angle out to any distance including, but not limited, to the outer lines.

In keeping with [11], *w_ij_* ∊ ℝ. That is, we allow all synaptic weights to take on both positive and negative values and, indeed, for weights emanating from a single input neuron to take on both positive and negative values depending on their target neuron, completely ignoring Dale’s Law. This is a common computational conceit, to aid mathematical analysis. Limitations imposed via Dale’s Law and a sign on weights certainly do influence computation but can usually be worked around by reference to larger population encodings, compensation between competing pathways, and offsetting everything from 0 to avoid such hard bounds. We have also omitted the hard bounds on weights defined symmetrically around 0 in [11], largely as these were never attained in the simulations presented in [11], but also, to aid the analysis.

We have just two neurons in our output layer. As in Herzog et al. [11] each output neuron computes a weighted sum of their inputs and adds independent noise,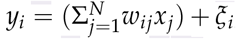, for *N* input neurons. As with [11], we allow for negative firing rates. Since weights are unbounded, firing rates are also technically unbounded. This greatly simplifies the analysis, but does lead to some open questions about whether natural limits on firing rates may provide natural limits on the results of an unsupervised bias. It has been suggested (*personal communication - WG*) that the network described in [11] might have been better served with some form of softmax function on the output layer. While this would definitely limit some runaway effects it is unlikely to qualitatively change the impact of the unsupervised bias, rather it would work as a learning rate stabiliser but the outcome would be the same. Issues such as these will be tackled in **Section 4.1**.

The goal of our decision network is to correctly distinguish a left input (*φ* < 0) from a right input (*φ* > 0). We have already moved our input encoding to be 0-centred, with a positive input representation for ‘right’ and a negative input representation for ‘left’. Therefore the network must develop positive weights from input layer to the ‘right’ output neuron and negative weights to the ‘left’ output neuron. The classification works by comparing the two output neuron firing rates and choosing the greater (more positive) firing rate, via a winner-takes-all mechanism. We have a more advanced analysis (in **Section 3.2**), which omits the winner-takes-all mechanism and arrives at qualitatively the same results as the current network. The main effect of the winner-takes-all mechanism is to separate the pathways more cleanly, leading to faster learning when learning is possible.

The final simplification which we can make to our network is a reduction in the number of input neurons. Herzog et al. typically used 50 input neurons per bisection task. This provides numerical stability and a sense of realism. For the mathematical analysis, given the task representation which we’ve described in the lines above, we need only a single input neuron per bisection task (here bisection task refers to a stimulus presentation with a given outer line separation, regardless of where the central line is projected). We do not need a second input neuron with an opposite slope on the tuning curves, although that would greatly aid in the numerical implementation of a more realistic system. So, for a given outer line separation we have a single input neuron, which drives just two output neurons via two weights (**Figure 3**). When we present with a different outer line separation, we assume that we use a different input neuron and a different set of weights; at that point it is irrelevant whether the output neurons are the same for each task or different per task, only their ability to reliably signal the choice direction matters. We have now arrived at the simplest network possible for classifying left from right input signals, while remaining compatible with the chosen reward learning scheme.

**Figure 3:**
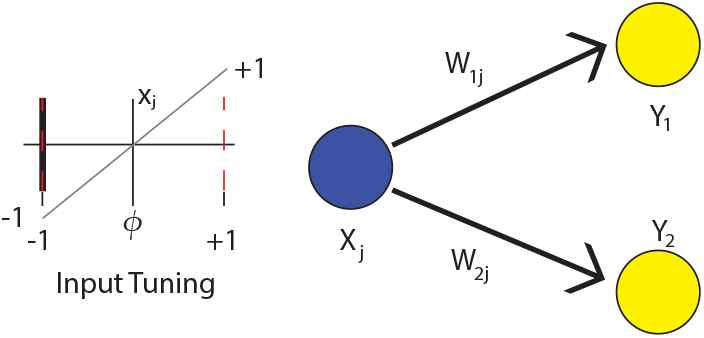
Simplest form of network which may peform a single bisection task. A single input neuron represents left of centre inputs as −1 and right of centre inputs as +1. The network must learn to associate output neuron 1 with the left input and output neuron 2 with the right input. Separate tasks are performed by similar but non-overlapping networks.

Now that we have described the network, we must clarify whether our use of *a* = 0 in the input layer obviates the unsupervised bias, since the input description now has expectation 〈*x_j_*〉 = 0. For a range of situations, including the correct solution, we could expect such a situation to nullify the unsupervised bias term via, 〈*e_ij_*〉 = 〈*x_ij_*〉 = 〈*x_j_* · *y_i_*〉 = 0, where *y_i_* is the output layer firing rate. We avoid this situation, largely via our task encoding, by using a desire towards negative weights on negative inputs in order to obtain a correct classification. In brief, since the unsupervised bias is defined *per synapse* we see for our linear network that,

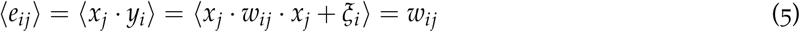

since 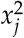 is always positive and 〈*ξ_i_*〉 = 0. We ignore the case *w_ij_*; = 0 as it is unstable and will immediately disappear in the long run average. More importantly, since each synapse can only exist in exactly *one* of the states above, we can confirm that we are *not* accidentally removing our unsupervised bias by moving the base value of *x* in our simplification of the classification problem. To be more precise about the value of 〈*e*_ij_〉 we further point out, that this formulation is before winner-takes-all is applied, after which the true value of 〈*e_ij_*〉 will lie between 0 and the values indicated above, in proportion to how many times that particular output neuron is active. Finally, we point out that this scheme expands to large numbers of input neurons targeting the same output neuron; what matters is the relationship between an *individual* input neuron and the respective output neuron. Our +/ − 1 input, with a crossover at the true bisector, encoding scheme combined with linear weights ensures that opposite inputs result in the same encoding in the output neuron. The value of the eligibility trace is dictated by the sign of *w_ij_*, which is stable across trials. In the case of multiple input neurons, this result extends across all of the input neurons and since we are looking at a stable average, even if the neuron of interest is ‘voting’ against the majority of his fellow input neurons, it will consistently do so across all trials. In short, our analysis does not fall foul of this potential pitfall hidden within the unsupervised bias hypothesis.

The merits of reward learning as a modelling paradigm, for this task, are discussed in **Section 4.2**. For the analysis we accept it as a given. Since this paradigm is based on updating weights in accordance with the difference between the predicted reward and the observed reward, in keeping with [11], we define *S*(*R*) = (*R* – 〈*R*〉). Reward is defined as *R* ∊ { − 1,1}, for respectively a failure/success trial, and 〈*R*〉 is the expected value of reward over trials and time. The fact that *S*(*R*) is defined in terms of both *R* and 〈*R*〉 conveniently allows us to maintain both *R* and 〈*R*〉 on the interval [−1,1]. This does not result in any loss of generality. Two alternative conditions can quickly be dismissed: if we extend, or restrict, this range we see only a modification of the learning rate (step size), 〈*R*〉 will track *R* no matter what the range; similarly if we make the range non-symmetric around 0, we will see an asymmetric biasing in the *R* values, in either the positive or negative direction, but 〈*R*〉 will still track this range accounting for the true average reward signal accurately. Updates, via *S*(*R*), will still be symmetric around this value. We define *S*(*R*) in keeping with [11], but we will not necessarily need to use a running average of reward as a proxy for 〈*R*〉, rather we will rely on the true task reward averages, 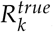 as will be described in **Section 3.1**

A summary of the simplified model equations is presented in Table 1. We will proceed to use them for the model analysis in **Section 3.1**.

**Table 1:** Simplified model equations. These are sufficient to perform an analysis of the original unsupervisd bias hypothesis of [11].

|  |  |
| --- | --- |
| Input representation: | $\phi = -1, \text{ left}; \phi = +1, \text{ right. } x(\phi) = \phi$ |
| Output representation, classification: | $y_i = x_j \cdot w_{ij} + \xi_i, \text{ winner-takes-all}$ |
| Weight representation: | $w_{ij} \in \mathbb{R}$ |
| Success signal (RPE): | $S(R) = (R - \langle R \rangle)$ |
| Weight update rule: | $\Delta w_{ij} = x_j \cdot y_i \cdot S(R)$ |

## 3. Results

### 3.1. Analysis of Unsupervised Bias hypothesis

We can now proceed with our analysis of the unsupervised bias as an explanation for failure to learn during task roving, initally from the perspective of the Herzog et al. (2012) [11] model, but subsequently from the point-of-view that no model could both (i) correctly model roving, and (ii) fail in its learning due to the unsupervised bias induced via an incorrect reward prediction error.

**Figure 4:**
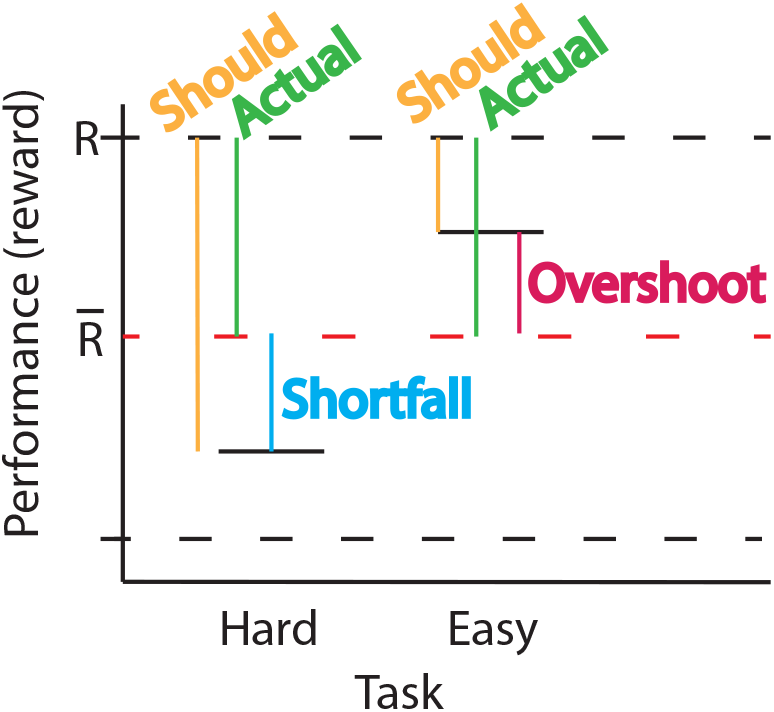
Schema for how an error in estimating a reward prediction average can lead to an overshoot or a shortfall in the reward prediction error. Overshoot corresponds to *ε*_1_ in the analysis, and shortfall to *ε*_2_.

Let us begin by (re-)defining the weight update equations:

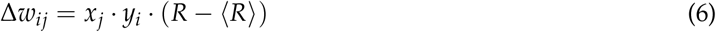

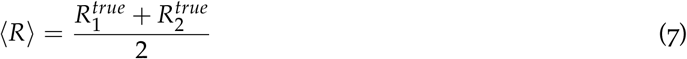

where 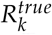 is the true probability of performing task *k* correctly. In our first analysis, we will look at the roving of two tasks of differing outer line distances, henceforth referred to as an ‘**easy**’ task and a ‘**hard**’ task. In keeping with the experimental methodologies presented in [22, 29, 11] we will assume equal ratios of task presentation, the advanced mathematical analysis (**Section 3.2** and **Appendix B**) can cope with non-equal presentation ratios. The results presented in [11] claimed an inability to learn to improve on either task, while randomly alternating the two tasks. Without loss of generality, we will denote task 1 the ‘**easy**’ task, and task 2 the ‘**hard**’ task. This means that,

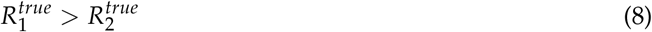

Note that, without the assumption of non-equal performance then no unsupervised bias can exist. We further define two ancillary variables which will aid greatly with the analysis,

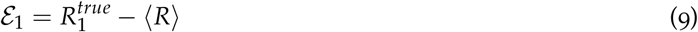

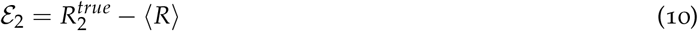

It is trivial to note that,

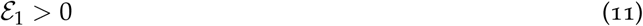

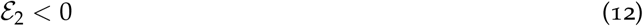

Let us now look, in detail, at the calculation of one of the weight updates. When *φ* = −1 is the presented stimulus, on task 1 (**easy**), and we correctly choose Left, we obtain,

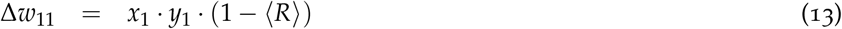

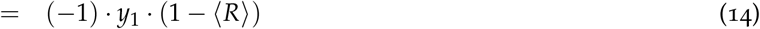

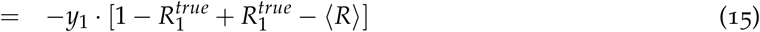

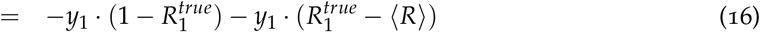

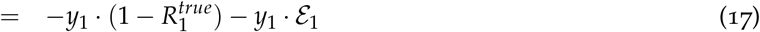

Following the same general form of this calculation, we can now fill out, in table form, the weight updates as they would be performed upon each potential trial outcome (Table 2). On any given trial, we have only one input presented to the network and one output option chosen, the winner-takes-all mechanism disables all other weight updates, so only a single entry from the table is active. It remains to be shown, whether the unsupervised bias which is exclusively represented by the terms *ε*_1_ and *ε*_2_ can ever actually prevent learning from occurring.

**Table 2:**
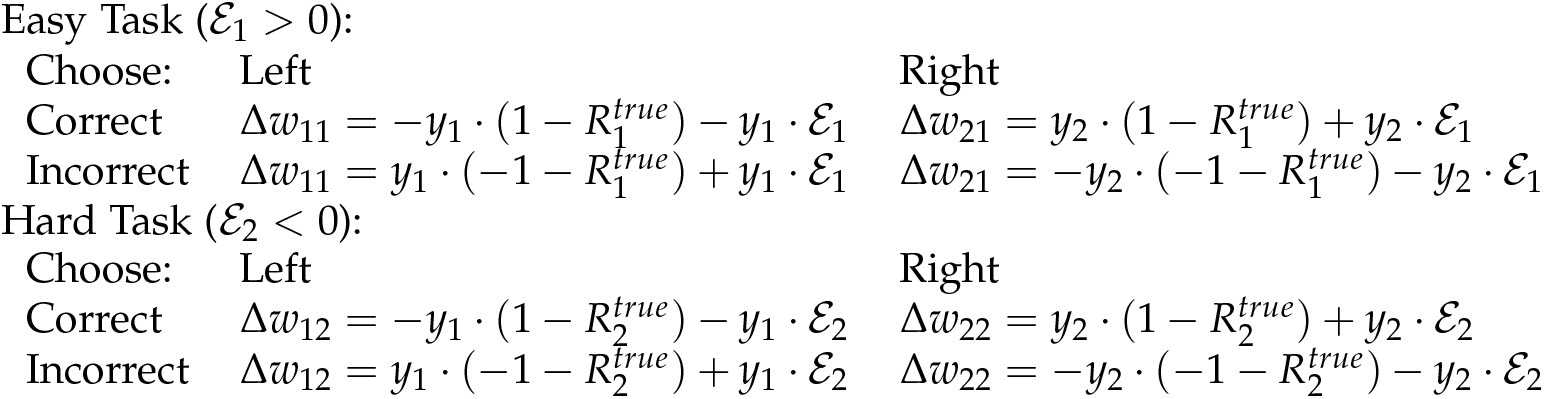
Weight update rules, per task and per trial outcome, in a non-overlapping task representation scenario. That is, the analysis of the simplified model of the unsupervised bias hypothesis.

|  |  |  |
| --- | --- | --- |
| Choose: | Left | Right |
| Correct | $\Delta w_{11} = -y_1 \cdot (1 - R_1^{true}) - y_1 \cdot \mathcal{E}_1$ | $\Delta w_{21} = y_2 \cdot (1 - R_1^{true}) + y_2 \cdot \mathcal{E}_1$ |
| Incorrect | $\Delta w_{11} = y_1 \cdot (-1 - R_1^{true}) + y_1 \cdot \mathcal{E}_1$ | $\Delta w_{21} = -y_2 \cdot (-1 - R_1^{true}) - y_2 \cdot \mathcal{E}_1$ |
| Choose: | Left | Right |
| Correct | $\Delta w_{12} = -y_1 \cdot (1 - R_2^{true}) - y_1 \cdot \mathcal{E}_2$ | $\Delta w_{22} = y_2 \cdot (1 - R_2^{true}) + y_2 \cdot \mathcal{E}_2$ |
| Incorrect | $\Delta w_{12} = y_1 \cdot (-1 - R_2^{true}) + y_1 \cdot \mathcal{E}_2$ | $\Delta w_{22} = -y_2 \cdot (-1 - R_2^{true}) - y_2 \cdot \mathcal{E}_2$ |

In order for learning of the ‘**easy**’ task to work we observe the following,

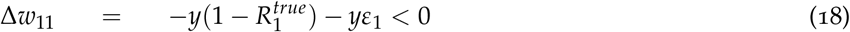

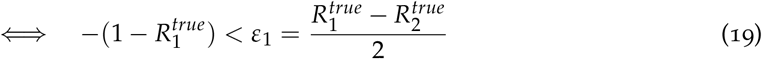

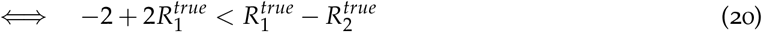

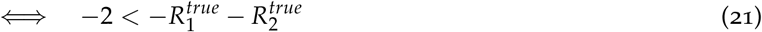

The condition Δ*w*_11_ < 0 is by construction of the classification task; negative weights are required in order to correctly classify a negative input as ‘Left’. The final line always holds, 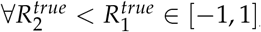, which is the basic setup of our scheme.

Similarly, for positive valued inputs, which must be classified as ‘Right’, we have the condition,

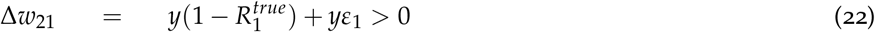

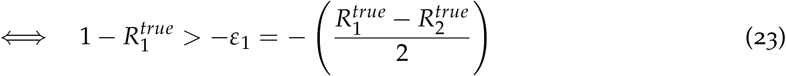

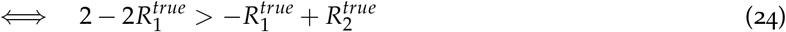

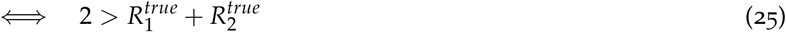

The final line always holds, 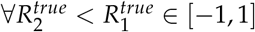.

Similarly on the failure trials, we need the same sign for Δ*w*_11_ and Δ*W*_21_ but now with the different equations, giving us,

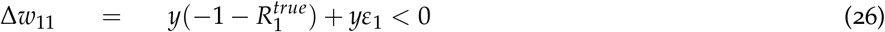

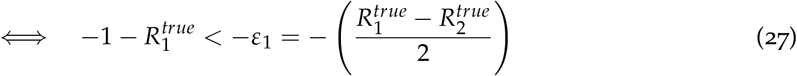

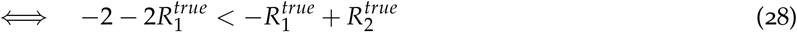

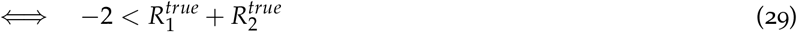

and,

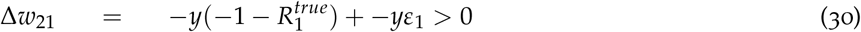

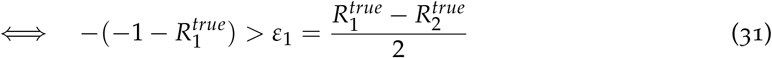

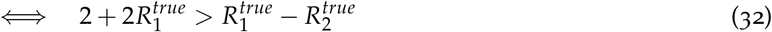

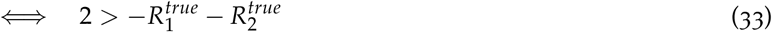

Simple inspection allows us to see that, 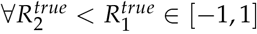, meaning that in all cases of presentation of the ‘**easy**’ task we observe *net* weight movements in the direction towards a correct solution.

It is similarly possible to carry out this analysis on the synaptic update rules for the ‘**hard**’ task. The details are presented in **Appendix A**. As with the ‘**easy**’ task, we find that in all cases an unsupervised bias from the other (roved) task can never be large enough to prevent learning.

This analysis, while surprisingly simple, is actually quite profound. What it means is that, in general, given a learning bias, inherent in the synaptic update rule, from a task which shares no neuronal overlap in representation, you can always expect the system to learn. What is not shown, but which follows quite easily, is that the unsupervised bias can only ever have two effects on non-overlapping tasks, (i) it can change the rate of approach towards the correct solution, accelerating or decelerating it, and (ii) it can overshoot the correct solution, leading to inaccuracy, as long as the bias is still present in the system. This result suggests that the conceptual understanding of the unsupervised bias hypothesis, as it applies to roving, cannot hold.

### 3.2. So how does the Herzog et al. network fail to learn?

The network simulations presented in [11] contained an unsupervised bias, by construction. They also strongly resembled the mathematical setup of our simplified model, analysed in **Section 3.1**. And they failed to learn in a task roving scenario. We will now show how this happened. In **Section 3.2.1** we will extend the mathematics of our simple model to show that the mechanism of failure in [11] is actually a side-effect of runaway postsynaptic firing rates. While we will conclude that this is an unsatisfactory explanation for roving, this does uncover a different aspect of the unsupervised bias. It turns out that the unsupervised bias has a far greater influence on shared neuronal task representations, than on nonoverlapping task representations. In fact, it is important to separate the effects of unsupervised bias into shared and non-overlapping task representation terms. We will numerically explore the separation of the two forms of unsupervised bias, and their implications, in **Section 3.2.2**.

#### 3.2.1 Analysis of left-right bias

We extend the analysis of **Section 3.1** in two manners. First, we recognise that correctly distinguishing a ‘left’ stimulus, or a ‘right’ stimulus, is already a sub-task on a given input pathway, and our performance on the two may not be equal. This gives rise to a left-right unsupervised bias on a given task. Then, we will point out that learning to correctly identify one sub-task on a given input pathway necessarily modifies performance also on the other sub-task. To account for this in our analysis we will need to slightly modify the encoding of our analytical network, giving all firing rates a positive offset from zero and adding a second input neuron. While the simplified model described in **Section 2.1.2** was sufficient for the analysis of an unsupervised bias induced from the other (roved) task, without these modifications it would not be sufficient for the analysis of a left-right unsupervised bias.

We begin by re-defining the error in the estimation of the reward prediction error in terms of expected reward,

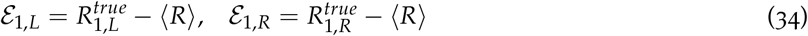

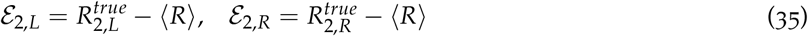

So, for example, *ε*_1,*L*_ is the error in reward prediction, or estimation, on the ‘easy’ task, when a left stimulus is presented. It is defined as the difference between the true expectation of performance on that task and the single critic average, (R). The subscript 2 refers to the ‘hard’ task and 〈*R*〉 to a right of true bisector stimulus.

We note in passing that *ε*_1_ and *ε*_2_ may be algebraically composed of the elements of Equations (34, 35). As before, we can assume that *ε*_1_ > 0 and *ε*_2_ < 0. Further, without loss of generality, we can assume that *ε*_1,*L*_ > *ε*_1,*R*_ and *ε*_1,*L*_ > *ε*_1,*R*_. Finally, as a starting point, we will assume that performance on the left-right subtasks is closer to performance as a whole on that task, than to performance on the other task. This means that,

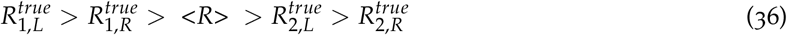

This will not hold at the end of the analysis, but it is a natural initial condition of any system, using reward learning, expected to correctly learn to distinguish left from right on two separate pathways. It also corresponds to the initialisation conditions of the network presented in [11].

We can now update Table 2, using our sub-task notation, giving us Table 3. Note that it is relatively simple to do this. Despite the fact that we have considerably more granularity in describing our potential errors in the update rule, the actual mechanisms in both cases remain the same, meaning our table has exactly the same number of entries as before. On the *Incorrect* lines of the table an interesting notational decision needs to be made, we can annotate the equation in terms of the stimulus actually presented or in terms of the incorrectly identified stimulus. Both are mathematically accurate and appropriate in different contexts. In the current context, we find it more appropriate to represent the bias in terms of the actually presented stimulus.

**Table 3:**
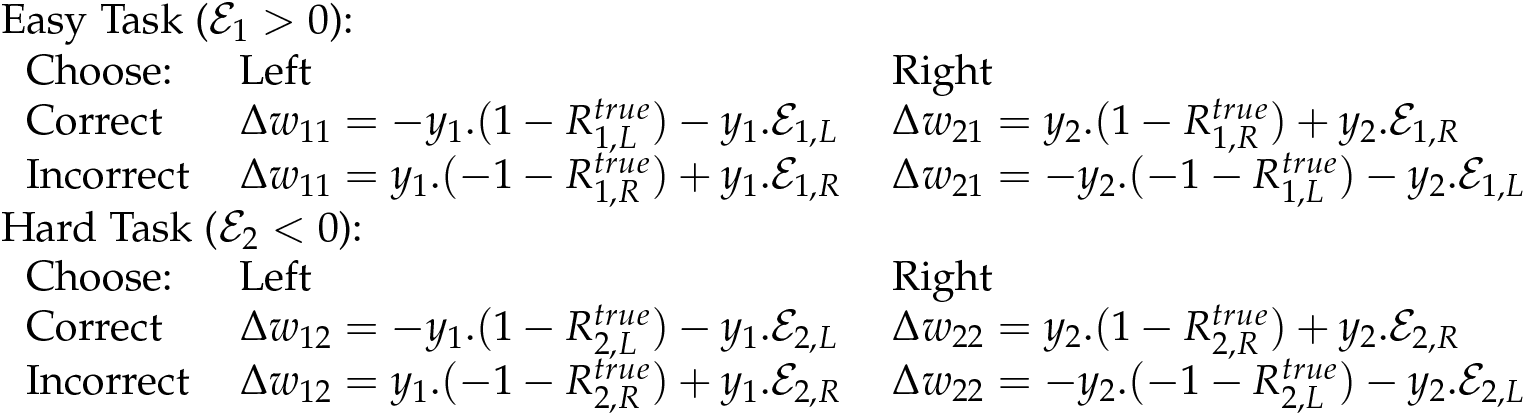
Weight update rules, per task and per trial outcome, in a non-overlapping task representation scenario. That is, where inputs on Easy Task are carried via separate synapses from inputs on Hard Task. In this case, we have more granularity in the left-right error in reward prediction than in **Section 3.1**. At this point we have not yet included supplementary input neurons.

**Figure 5:**
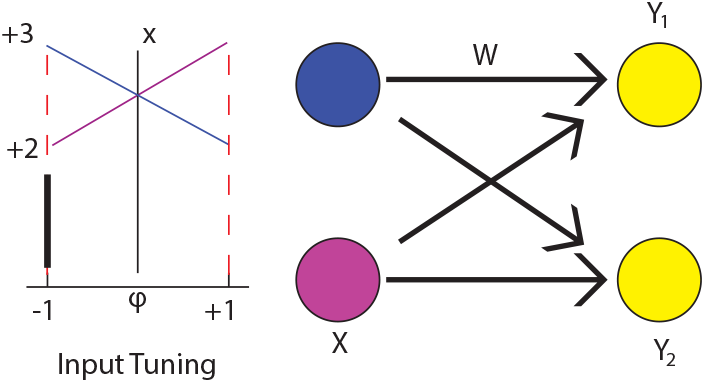
Network relevant to performing a single bisection task, where task encoding is accounted for in the analysis. In order to analyse where the Herzog et al. network [11] really fails to learn we need a second input neuron. Both input layer neurons have only positive firing rates, but opposite tuning curves. Separate tasks are performed on similar but non-overlapping networks.

At this point, we need to modify the task encoding from **Section 2.1.2** slightly. While the simplified encoding was sufficient for comparing two non-overlapping task representations (**Section 3.1**) it cannot be used here for two reasons, (i) modifying weight Wtt for an input of *φ* = 1 will modify our classification, also, for *φ* = −1, and we don’t yet have a mechanism in the simplified analysis for dealing with this, and (ii) actually the correct classification boundary *W* > 0 comes about accidentally here and will always be learned, no amount of bias could ever overcome it since it involves a sign change. We need to design a system, for analysis, in which a systemic weight bias could potentially interfere with learning. So we restore aspects of the full Herzog et al. network encoding [11], although still greatly simplified. Instead of 50 input neurons per task, we can imagine instead that it has only two input neurons. Furthermore, we can fix these neurons such that one input neuron has a firing rate of 2/sec for an input of *φ* = −1 and 3/sec for an input *φ* = +1, meanwhile the encoding of the other input neuron is the reverse of this. This is enough to provide an intuitive understanding of what happens in the Herzog et al. [11] network. The understanding is of course supported by simulations, and will be further reinforce by a numerical analysis in **Section 3.2.2**. The portion of the new network devoted to a single bisection task is depicted in **Figure 5**.

The change in encoding enforces a slightly more difficult balance of weights in order to find a solution, hence its susceptibility to unsupervised biases. Weights from the input neuron with positively sloped tuning function should provide strong drive to output neuron 2 (the ‘right’ choice neuron) and weaker, or ideally no, drive to output neuron 1. For the other input neuron this is reversed. Since winner-takes-all still simplifies the output encoding, and hence the weight update assignment, we see that for a given trial the weight update equation from the lower firing input neuron to the output neuron will move in the same direction as that of the higher firing input neuron, but it will be slightly smaller in magnitude leading to a separation upon repetitions. What is important to retain from this is that, (i) the weight update rules are still doing the correct thing, and (ii) the new encoding is only to avoid logical fallacies in the analysis and does lead to a separation in choices over time.

With this new encoding, with multiple input neurons per task, we find it useful to simplify Table 3 considerably as follows:

**Table 4:** Markup of weight changes observed in Table 3. This simplified notation helps with analysis when there are multiple input neurons per task. Weight update equations are split into two terms, which may have positive, +, negative, −, or initially positive but later negative, ⊕, effects on the respective synaptic weight value. Only one weight update rule is performed on a given task. The probability of it being a given update rule is given by *Prob(chosen direction|presented direction*, *task).* Given the initial conditions described by Equation (36) the easy task has a natural ordering from upper-left, clockwise, to lower-left. The left-right imbalance in both overshoot-drag and in frequency of occurrence leads to a run-away process, whereby eventually ‘Left’ is chosen all of the time.

|  |  |  |
| --- | --- | --- |
| Easy Task ($\mathcal{E}_1 > 0$): | | |
| Choose: | Left | Right |
| Correct | — — | + $\oplus$ |
| Incorrect | — $\oplus$ | + — |
| Hard Task ($\mathcal{E}_2 < 0$): | | |
| Choose: | Left | Right |
| Correct | — $\oplus$ | + — |
| Incorrect | — — | + $\oplus$ |

We look at each equation in Table 3 and observe that for each entry there are two parts to the weight update equation, the pure learning term (e.g. (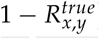)) and the unsupervised bias term (±*ε_x,y_*). We now markup each entry in Table 4 with whether the respective equation component can be expected to increase, +, or decrease, −, the relevant weight. The circled entries, e.g. ⊕, begin as the entry inside the symbol but can change sign when the ordering of sub-task performances defined in Equation (36) no longer holds. This schema allows us to avoid a doubling of the number of equations in Table 3 to account for the doubling in the number of input neurons. We are allowed to do this as the increase in the number of input neurons does not change the sign of, *y*, the post-synaptic firing rate and we are largely only interested in the sign of the weight updates. Also, the actual *y* values multiply by both terms in a given equation.

When the symbols in a table entry are in agreement, we will refer to this situation as *overshoot*, and when they differ, we will call it *drag*, referring to the effect of the unsupervised bias on the correct gradient (these correspond to the effects of Overshoot and Shortfall, respectively, in **Figure 4**). In explaining our analysis, we will make one further simplifying assumption, that we are performing better than 50% correct on all tasks. This will allow for the development of a simple intuitive understanding of the mechanisms involved in the unsupervised bias, but would lead to a tedious iteration of sub-cases if omitted. A full accounting has been checked and it corresponds with this subset presentation. The numerical solutions provided in **Section 3.2.2** will extend the analysis to cover all cases.

If we examine the **Easy** task equations (Table 4). The natural ordering of the table entries, based on the direct probabilities of correctly or incorrectly classifying each stimulus is that the upper-left entry occurs most often and the frequency decreases in an entry-by-entry clockwise manner around the other table entries. We identify immediately that for our most frequently experienced weight update (the upper-left entry) we experience an ongoing *overshoot* stemming from −*ε*_l,*L*_ in the original equations. Furthermore, this is under-compensated for by the *drag* in the least commonly visited weight update (the lower-left entry). In fact, if the performance of left and right ever get so separated that right now under-performs 〈*R*〉 then even the *drag* on this equation will disappear. This column looks like a runaway process.

By contrast, the right-hand column (containing the weight changes occurring whenever the right-hand output neuron is correctly or incorrectly selected) will observe quite mixed conditions. On incorrect trials we see considerable *drag* from the highly performing *ε*_l,*L*_ term but this is balanced by a deep potential well (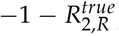). On the other hand, the more frequently experienced correct trials lead to an initial *overshoot* in learning.

**Figure 6:**
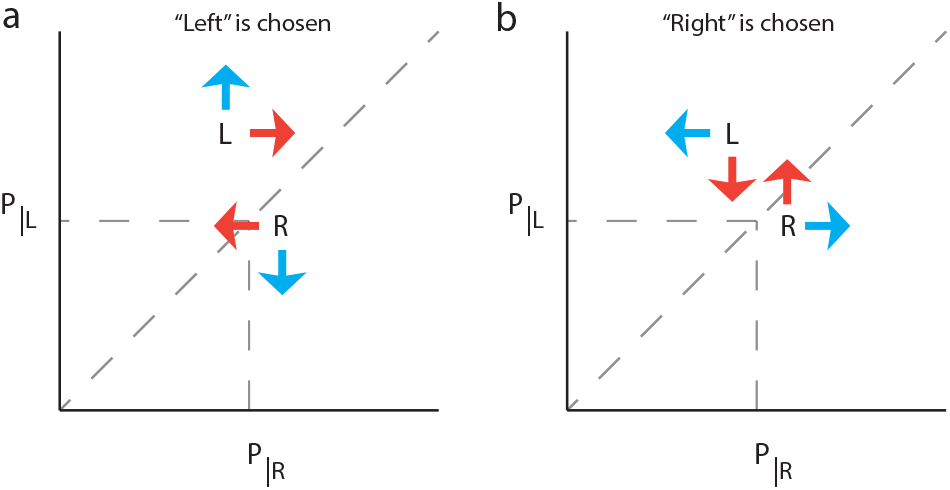
Sketch of the probability of choosing “Left” (L) or “Right” (R) given an input which is either left or right of the true bisector, under the initial conditions described by Equation (36). Vector arrows show the impact of choosing Left (**a**) or of choosing Right (**b**). Due to the overlap in task representations, choosing Left when a left stimulus is presented not only decreases the likelihood of choosing Right the next time that left is presented (**a, cyan** vectors) it also decreases the likelihood of choosing Right even if right is presented (**a, red** vectors). The converse is the case if Right is chosen (**b**). This overlap in the task representations, combined with an initial imbalance in how frequently each scenario will arise, leads to a run-away system in which Left is eventually always chosen.

It turns out that the magnitude of the weight update on the correct trials for both task outcomes is initially effectively the same (ignoring for a moment the effects of the post-synaptic firing rate *y*). This, despite the new notation introduced, is since all weight updates are actually with respect to 〈*R*〉 and is partly a result of initial conditions (Equation (36)). However, the system visits the upper-left element of the Tables of weight updates more frequently than the upper-right element, and vice-versa for the lower row entries. This leads to slightly larger weight updates for ‘left’ tasks than for ‘right’ tasks.

Importantly, since ‘left’ and ‘right’ share neuronal representation, by altering the probability of choosing ‘right’ for a future ‘left’ stimulus, we are also altering the probability for a future ‘right’ stimulus. This is schematised in **Figure 6** (red vectors are undesirable impacts on future choice probabilities). The imbalance in frequencies, combined with the further relative infrequencies of visiting the lower-row elements of the weight update tables and the ultimately disappearing *drag* on the lower-left update rule, leads to a run-away weight update rule. This eventually leads to the left-hand option effectively always being chosen. This effect is further enhanced by the inclusion of the magnitude of the post-synaptic firing rates, *y*, in the analysis.

The analytical approach taken in this section, equally extends to the **hard** task and is only reinforced by relaxation of the initialisation assumptions contained in Equation (36). Most importantly it highlights for us, three points: (i) correctly distinguishing left from right can be considered a task, and in **Section 3.2.2** we will show mathematically that this also contains a form of unsupervised bias; (ii) it is not possible to study the effect of synaptic weight biases without also taking into account task encoding (c.f. the red vector arrows in **Figure 6**); (iii) the actual method by which the Herzog et al. [11] simulation fails to learn is a run-away system. We have confirmed this last point by reproducing their simulations (the results look similar to the single task simulation in **Figure 9**). The method by which their model fails to learn during roving is by learning to always choose ‘Left’. While we will show, shortly, that this is the result of an unsupervised bias it is not a realistic explanation for failure to learn in perceptual learning. In **Section 3.2.2** we will move on to a numerical analysis inspired by the Herzog et al.[n] system. This will allow us to separately quantify the effects of the different components and to develop a more general theory of the unsupervised bias.

#### 3.2.2. Separation of the unsupervised bias into subterms

We switch now to a numerical analysis of the unsupervised bias. We have developed a mathematical description of a perceptual learning system for three different output encoding possibilities, (i) the winner-takes-all schema presented in Tables 2 and 3, (ii) a linear output encoding scheme in which the task encoding is effectively the same as above, but without winner-takes-all, and (iii) a binary output encoding scheme, which emulates the winner-takes-all but removes the ability to have a truly run-away system since firing rates are either 1 or 0. For our analysis, we will focus on the results from the linear output encoding scheme, the results of the first (WTA) encoding scheme should be close to identical but require technical decisions to be made about firing rates which are beyond the reasonable scope of this document. Binary output encodings are mentioned briefly in **Section 4.1**.

The mathematics of the linear output encoding analysis can be derived from first principles (**Appendix B**) but they can be sumarised as follows. The likelihood of choosing correctly is based upon the separation in the expected firing rates of the output neurons, an increase in this separation will lead to a greater likelihood in future of correctly classifying the same inputs. However there are two sources of bias which may corrupt this correct learning path. As we saw above, there can be a bias from a task with an over-lapping neuronal representation (**Section 3.2.1**) and separately from a task with a non over-lapping neuronal representation (**Section 3.1**). This gives us an equation with three major, related, parts. Firstly, we have the main gradient of learning, based on the true performance on that task (similar to the (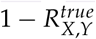) in **Section 3.2.1**). This affects not just future outcomes on the currently presented task, but also outcomes on tasks which share some neuronal representation with the current task (via shared weights; this is a component of the red vectors in **Figure 6**). Secondly, we have the influence of the unsupervised bias where we have under- or over-estimated our performance on the current task. This bias also has knock-on effects via the shared neuronal representation, changing the weights, on other tasks as well as affecting future performance on the current task. And finally, we have the effects of the unsupervised bias coming from tasks which do not share in the task representation of the current task. These influence the magnitude of our unsupervised bias as it is experienced on the current task, which again will have a downstream effect on other tasks which share in the current neuronal representation. We refer to the first term as pure learing, the second as internal bias and the third as external bias. Their separation is important as only the internal bias has a direct effect on its own future value, via the shared neuronal representation between tasks. See Equation (B.28) for a mathematical expression for these terms.

A simple visual example will hopefully make the influence of the separate components clear. **Figure 7** shows numerically calculated vector flow fields for the performance when a ‘left’ stimulus is presented and a ‘right’ stimulus is presented as a given *task* in an overlapping representation scenario. **Figure 8** extends this scenario to the case where there is a separate external/hidden task which does not share any neuronal representation overlap with the sub-tasks presented in the panels, but which contributes in equal proportion to the reward prediction and thus biases the learning in these observable tasks. Extensive simulations have verified the applicability of this approach to the Herzog et al. [11] network, showing that in the deterministic limit the simulations follow the flow lines of the vectors shown here. It should be noted that the derived flow fields for those simulations have a much flatter central region in all panels due to an extremely large *similarity*, or overlap, term in the neuronal representations of left and right.

**Figure 7:**
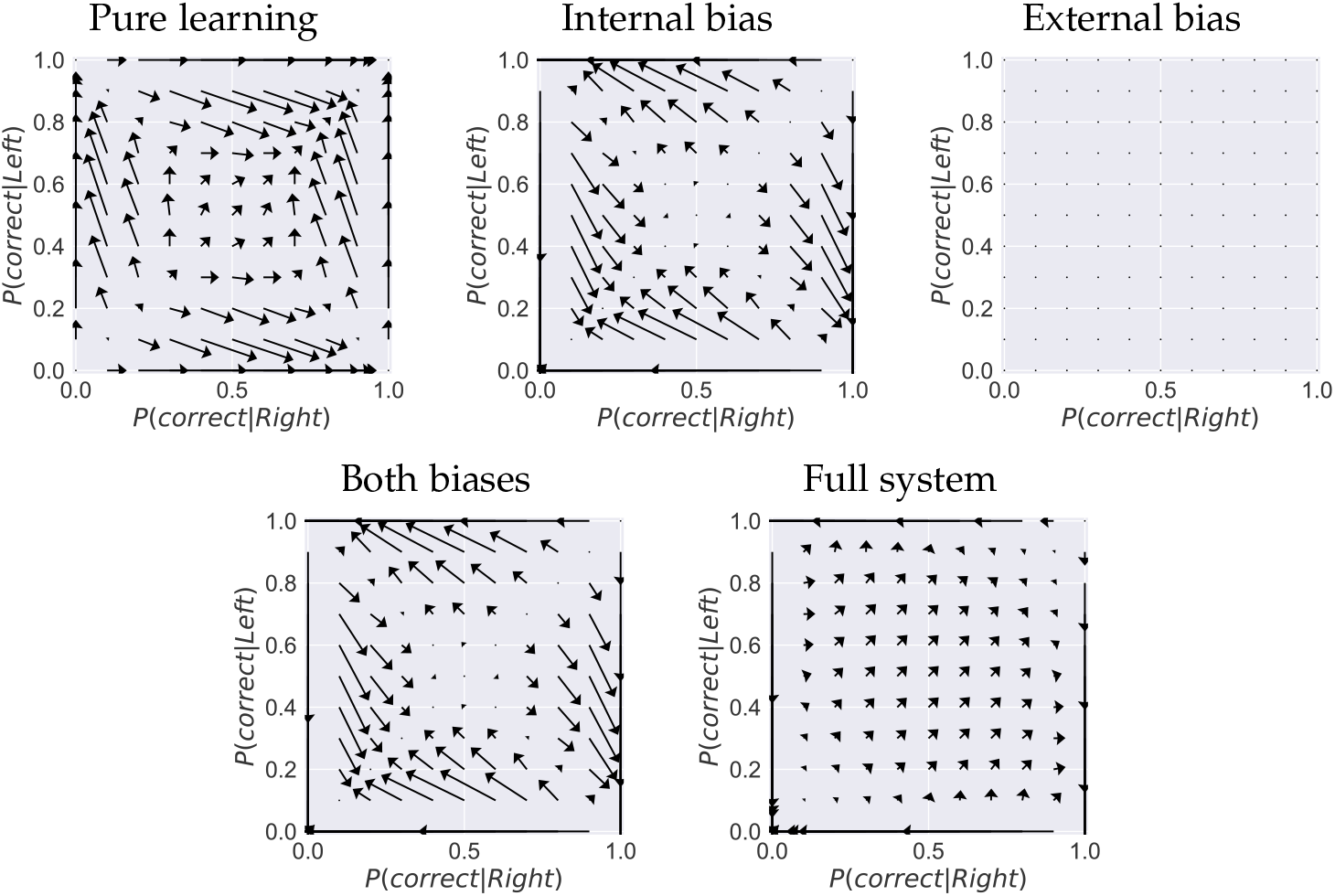
Vector fields for average effect of task performance on subsequent task performance on a simple left-right learning task with no external bias. Axes express probability of performing each task correctly given that a particular task has most recently been presented. Lengths of arrows are normalised across panels. Notice how the *biases* have a net detrimental effect on the upper and right hand borders, this is the run-away system where perfect performance on one task destroys performance on the other task over time. In the lower-right panel we see the combination of all of the effects in the update rule, the *unsupervised bias* effect outweighs the pure learning effects along the edges. For a system initialised with approximately equal performance on both left and right tasks, we may not expect this run-away behaviour to emerge as the natural path is along the diagonal to a region of zero flow close to perfect performance on both tasks. This is the case, for a limited region of parameter space, in the Herzog et al. [11] model on a single task. Once it has emerged however it is a relatively stable system. Perfect performance on one task and perfectly imperfect performance on the other is an unstable fixed point, but with a flat gradient in its immediate vicinity making it an effective attractor.

In the examples shown the **external bias** task has perfect performance in both left and right sub-tasks. This gives us a maximal possible unsupervised bias which can only be increased by changing the ratio of presentation of the tasks, presenting the hidden task much more frequently. The external bias is only able to change the lengths of the vectors in the **external bias** panels and not their orientation. Furthermore, it cannot affect the vectors in the **pure learning** and **internal bias** panels. Inspection of the effects of the biases suggests that the unsupervised bias can never truly counteract the **pure learning** term. Rather, the **internal bias** acting via overlapping task representations and a mechanism of run-away performance on a single sub-task is the most obvious mechanism for failure to learn in this system (as previously demonstrated in **Section 3.2.1**). An extreme example is provided in the simulations in **Figure 9**. There we see that the external bias is actually supporting learning and its absence leads to a failure to learn which is driven by the **internal bias**.

The discrepancy between our results and those demonstrated in [11] are due to their use of an interpolation function in their estimation of discrimination thresholds. When the network acquires the lopsided left-right behaviour, which we have demonstrated here, that function utilises the nearest data point which turns out to be a very poor estimate. We have avoided this problem by plotting proportion correct and probability correct in all cases. However, even in our case, if we attend to only the averaged left-right performance **Figure 9** (**green line**) it would be easy to overlook the unbalanced left-right behaviour.

In this section and **Section 3.2.1** we observe that the mechanism for failure to learn in a model similar to the one by Herzog et al. [11] is a run-away regime. This occurs when the **internal bias**, which is one aspect of the unsupervised bias, conspires with run-away firing rates to lead to a regime in which simulated subjects only choose one of either ‘left’ or ‘right’, no matter what the inputs. This has been confirmed first by simple mathematical logic of the weight update rules, then numerical investigation of the implications of performance on weight updates and, of course, detailed analysis of simulations. Such a mechanism of failure, while resulting from an unsupervised bias, is incompatible with explaining failure to learn during roving.

**Figure 8:**
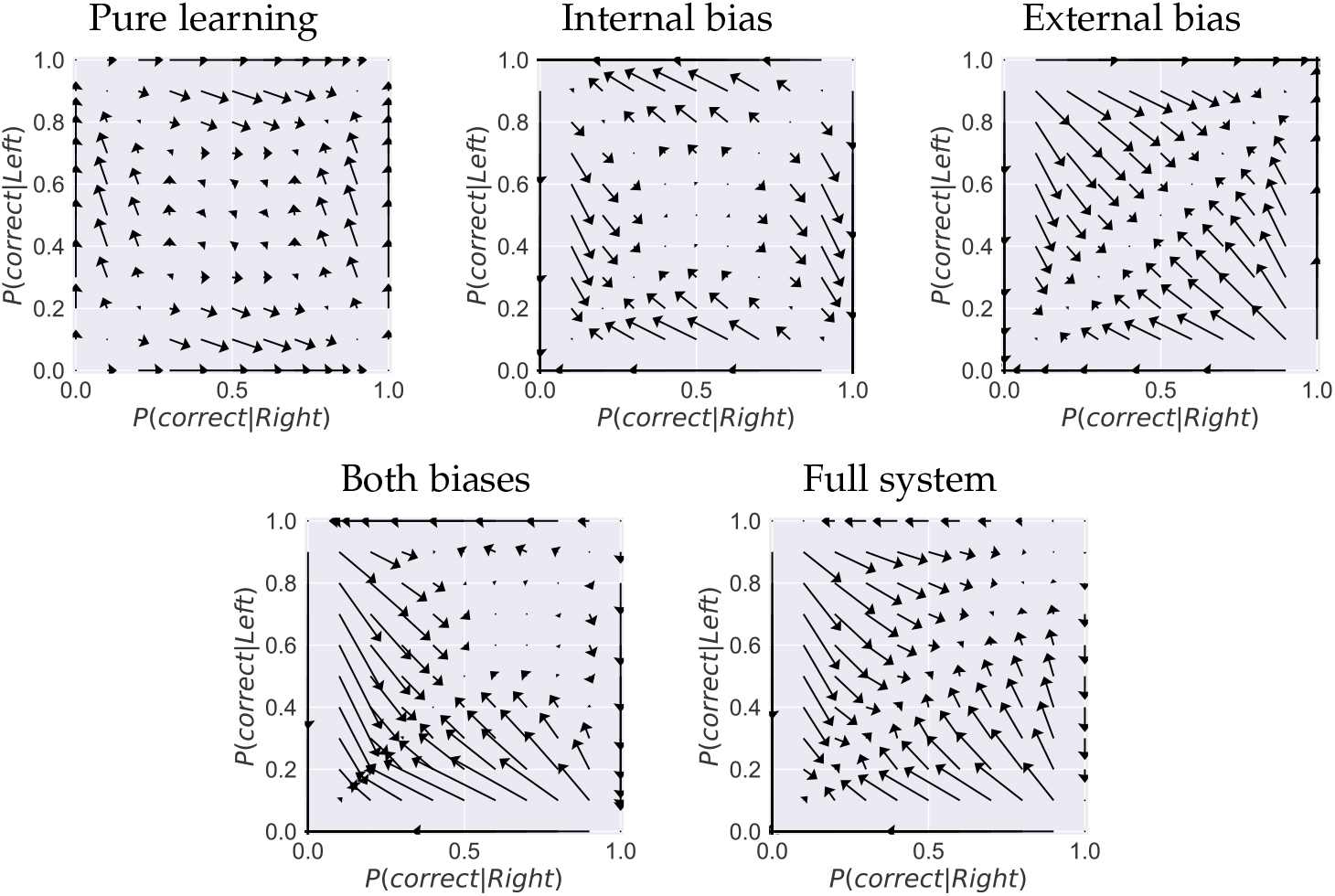
Vector fields for average effect of task performance on subsequent task performance on a simple left-right learning task with a hidden external bias from another left-right learning task with perfect performance. Axes express probability of performing each task correctly given that a particular task has most recently been presented. Lengths of arrows are normalised across panels, but must be doubled in length for comparisons with **Figure 7**. The main difference between the two figures is the presence now of a strong effect from the *external bias*. Less obvious is the fact that some vector angles are slightly different when compared with **Figure 7** due to the presence now of four sub-tasks rather than two. (The composition of *A* has changed). The most important point to note is that the external bias may only change the length of its related vectors, it may not change their angle. A negative bias will change make the arrow point in the opposite direction to that seen here. But our example uses the maximal value of external bias allowable in this system, meaning this is the largest effect it may have in the system.

More importantly, the directions of the vectors describing the impacts of the unsupervised bias on future performance appear to be approximately orthogonal to the direction required to truly prevent learning during task roving. Rather, they predict exactly the method of failure to learn observed in the simulations. At this point, the unsupervised bias hypothesis is largely incompatible with explaining roving.

## 4. Discussion

In **Section 3** we began with an analysis of the unsupervised bias hypothesis as it is presented in the paper by Herzog et al. [11]. We showed that the hypothesis, in its presented form, could not explain failure to learn during roving. We then expanded our analysis to demonstrate that the actual mechanism, of failure to learn, of the simulations by Herzog et al. [11] illustrates the necessity to bring neuronal task representation into any analysis of unsupervised bias. We did this, first analytically and then numerically, demonstrating an accurate accounting of the actual method of failure to learn, seen in the simulations. This analysis allowed us to highlight three forms of learning in a multi-task reward learning paradigm, which we called pure learning, internal bias and external bias. The external bias is the form of unsupervised bias hypothesised by [11] to explain roving. We showed that, in fact, this bias is more likely to have a stabilising effect on multi-task learning. Whereas, the internal bias, via shared neuronal task representations is the much more likely source of issues to do with learning in such a case.

**Figure 9:**
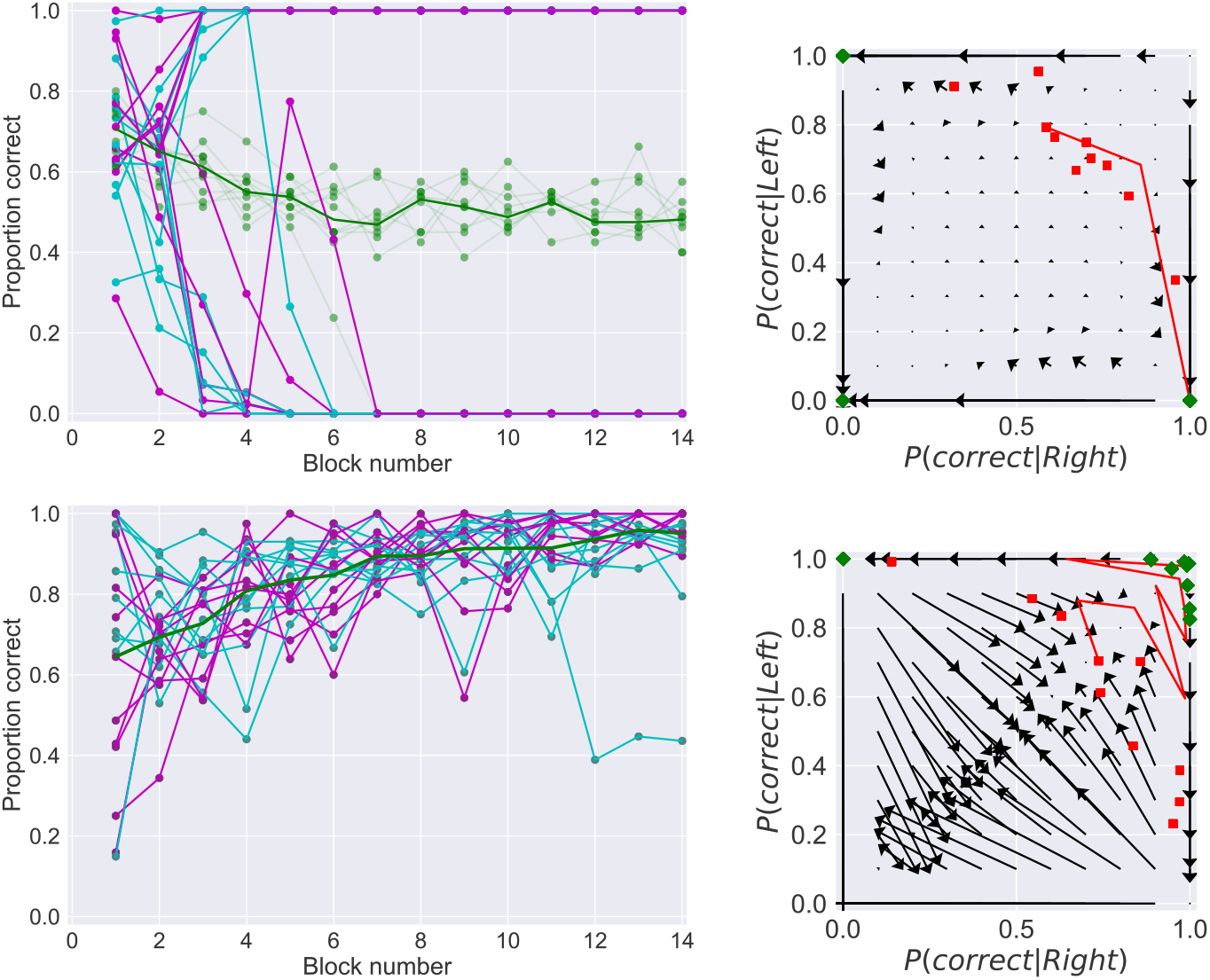
Top row: performance in a single task with no external/hidden bias. Bottom row: performance in a task with a strong external/hidden bias, which is always correct. Green lines are subject performances across left and right on a given task, cyan and magenta encode performance on left and right sub-tasks respectively. Note, for these simulations winner-takes-all has been disabled to more closely resemble the mathematics. Noise is also smaller than in [11], to smooth the trajectories and allow reproducible behaviour. In the vector plots, red squares denote initial performances of each subject, green diamonds their final performance. A single trajectory is depicted, in red, corresponding to the trajectory for the subject for whom the individualised flow field is derived. Counterintuitively, the mathematical prediction and observed simulation behaviour is more subject to failure to learn via the unsupervised bias in the case of only **internal bias** than in the case including a strong **external bias**. In the latter, the bias strongly *helps* performance.

In this section we will discuss alternative explanations for the failure to learn during roving. In **Section 4.1** we will examine whether our analysis in **Section 3** has been truly exhaustive. We present four alternative scenarios and examine whether any of these can combine with the unsupervised bias to induce a reliable failure to learn in a task roving scenario. Finding each solution implausible, we will discuss briefly alternative attempts at modelling perceptual learning and how they may be applied to roving (**Section 4.2**). Finally, in **Section 4.3** we will present an alternative hypothesis which may explain roving. In it, we will combine the single critic hypothesis, which originally lies behind the roving theory of [11], with recently published experimental results.

### 4.1. Can an unsupervised bias ever explain failure to learn during roving?

In **Section 3.1** we showed that the simplistic interpretation of the original unsupervised bias hypothesis [11] cannot explain failure to learn during roving. Furthermore, we have a mathematical analysis which explains the actual method of failure in simulations is an unexpected left-right unsupervised bias (**Section 3.2**). Due to its reliance on run-away processes, such as firing rates, this method of failure is far from realistic. Moreover, on a classification level, the left-right bias implies that failure to learn as seen during roving should be a much more widely seen phenomenon than we find in the literature. We turn now to examine whether there exists alternative task encodings which may lend themselves to the unsupervised bias hypothesis.

In total, we envisage four methods by which an unsupervised bias may impact on perceptual learning: (1) The pure, original, hypothesis that performance on one task leads directly via an over or underestimation of the average performance to mis-applied synaptic weight updates on the other task. (2) Via the shared neuronal representations examined in **Section 3.2**. (3) An expanded output encoding might allow for some reduction in the covariance term for the individual output neurons with respect to the task performance. This could then allow the bias terms to have greater impact relative to the pure learning term. (4) A higher-dimensional, recurrent, task encoding allows for a more complex interaction of bias with task performance. We will discuss each of these methods in this section.

(1) and (2) have been explored in detail in **Section 3**. (1) has been shown mathematically to be incompatible with explaining roving. At worst, it either slows learning or enforces an upper bound on performance. (2) has been shown to lead to run-away firing rates and an implausible method of failure. This could conceivably, however, be rescued by biological processes such as firing rate homeostasis. We have, separately (**Appendix C**), developed a numerical analysis for binary output layer firing. This strictly limits the separation in firing rates of the two output neurons to 1/sec. In such a case, the run-away edge cases disappear from the vector flow fields (not shown here). So then, the left-right inability to learn comes about at some separation of firing rates greater than 1/sec and less than infinity. In fact, our analysis shows that a firing rate separation of greater than 20/sec is already enough to lead to the run-away process shown in **Section 3.2**. This should not be considered a precise but rather indicative value, as it is dependent on the scale of the noise, which was not fitted to experimental data in the original Herzog et al. model [11].

We have pursued this idea of maintaining the separation of firing rates within certain limits, so that there is always a non-negligible probability that upon presentation of either stimulus (left or right) on a task, the system will choose either output. To date, in simulations, we have used neuronal intrinsic excitability and different forms of weight normalisation. The results were initially promising but the failure to learn during roving appears to be more accurately ascribed to a limitation in the ability of normalised weights to solve both tasks, an artefact of the system design. Further analysis, using entirely separate output neurons for the two outer line distance tasks, which leads to per task normalisation of the weights *does not* suffer from the unsupervised bias.

This can be explained by examining **Figures 7** and **8**. There are two sources of unsupervised bias in the figures, the internal bias and external bias. By examining closely Equation (B.28) we find that the external bias, that is the bias from the other task, can only ever change the length of the vectors in the upper-right panel in **Figure 8**. This is crucial to interpreting whether the unsupervised bias hypothesis can ever explain failure to learn during task roving, since it means that the *angles* of those vectors can not be modified. We have not derived a precise mathematical formulation for learning under firing firing rate homeostasis or weight normalisation. But we can say that these will largely change the *lengths* of the vectors around the edges and much less so their angles.

For the external bias to disrupt task performance it needs to be negative rather than positive (the example has a positive bias). This means that the external/hidden task must be performing with a probability of being correct less than 0.5. The perceptual learning experiments have been calibrated to have a much higher than 0.5 probability of correct classification. Further, the setup is symmetric so we would expect then, for that task which is performing so badly, to quickly learn, being helped by its own both internal and external biases, to ultimately remove this influence.

Again, it appears that the internal bias is the much more credible form of unsupervised bias which could, potentially, disrupt learning. It is possible to imagine a cycle in the vector fields which would never reach the borders, where one sub-task would perform correctly all of the time, but rather performance would be directed out towards such a border (over performance on a sub-task) before the influence of declining performance on the other sub-task would lead to a strong correction towards relatively balanced performances on both again. This is possible, but seems unlikely. If such a case were true then we would also expect to see its effects in learning a single task (**Figure 7**) and, of course, this is not the case.

(3) requires that the pure learning and, potentially also, the internal bias terms be *drowned out* so that the external bias signals in the system are so strong that they lead to a relatively *flat* vector field in the center of the performance diagram. Again, methods of normalisation or homeostasis may be required to keep this scenario biologically plausible. In the absence of the external bias this effect would not occur and normal learning could take place, but in its presence we might see a regime of zero learning for large ranges of performance values. The most obvious way to induce such a situation is by vastly increasing the relative presentation rate of the external/hidden task (i.e. not the one we’re plotting in our diagrams). This has already been tested in human subjects and appears to have no effect, learning occurs as normal on the over-presented task and the under-presented task has a learning rate relative to the number of times it is presented.

**Figure 10:**
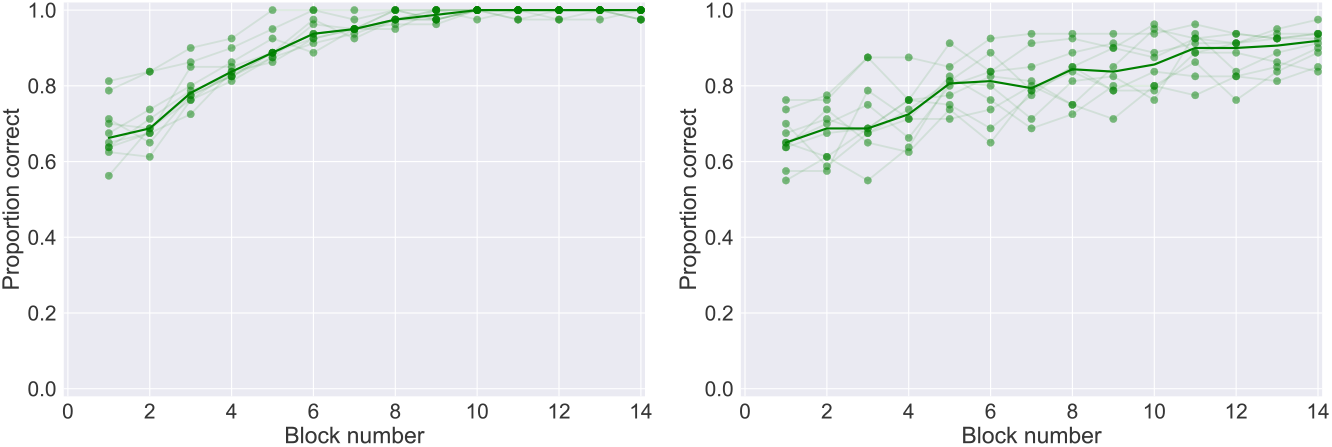
Effects of changing output population size on a simulation with 50:50 left-right cognitive biases. Demonstrating the effects of expanded output population on a simulation without this model addition is relatively pointless since in both cases the simulation doesn’t learn, but one is worse than the other. Here we show that the expanded output population slows the learning, but is otherwise irrelevant. Left is the original single output neuron per choice encoding, Right uses voting by 100 neurons per choice.

An alternative method by which we could ‘flatten’ the central performance regime of the **pure learning** term is by expanding the output population. In our main analysis we have stuck with just two output neurons and we’ve examined the effects of the unsupervised bias on the weights leading directly to these two neurons, and the subsequent effects of the synaptic changes on future task performance. It is entirely reasonable that an entire population of neurons are encoding a decision. If we then focus on a single representative neuron in each output population (‘left’ and ‘right’) we will see that sometimes the firing rate, on a given task, of this output neuron may be less correlated with the decision taken and consequently with the feedback received. Mathematically we can envisage this as a relative downweighting of the covariance term, relative to the biases, in Equation (4).

To test this, we have modified the simulations from [11] by mathematical expansion of the output population (**Appendix D**). This allows us to vary the output population size, still splitting it equally between ‘left’ and ‘right’ choices, while maintaining a constant signal-to-noise ratio. This means that we do not have to worry about having made the task easier or harder when we vary the population size. In this expanded output population encoding we assume that each neuron receives independent noise, which is typically the worst possible case for learning. The effect of the expanded output population encoding is most strongly observed in regimes where performance is approximately 0.5. This is the point at which the probability that our individual output neuron is correlated with the decision taken, and subsequent reward/feedback signal received, will be lowest.

We have implemented this system in our working model of perceptual learning and failure to learn during roving (**Appendix E**) and it appears to work extremely well. On blocks where only one task type is presented, the system correctly learns to distinguish left from right. Whereas on blocks where two tasks, of differing difficulties (important to maintain an unsupervised bias), are presented we see some degree of disruption of learning due to this mechanism. A full examination of this model will be presented in a later paper, but the basic result relevant here is that while the expanded output population does clearly dilute the effects of the **pure learning** term it requires an interaction also with phenomena such as weight normalisation to fully prevent learning in a stimulus roving paradigm. The expanded output population alone only slows learning, and even then only somewhat, as one would expect from any mechanism which dilutes an associative learning rule.

A final comment is necessary regarding this expansion of the output population as a method for tweaking the unsupervised bias influence in our equations. While this modification of the system does somewhat inhibit learning and while it technically is as a result of changing the balance of **pure learning** and **bias** terms in Equation (4), it is hard to equate this effect with the intent of the unsupervised bias hypothesis. It is rather a generic dilution of learning and affects both roved and non-roved scenarios. This is most evident in **Figure 10**, where expansion of the output population slows learning in a non-roved case. It has, of course, greater effect on the roved scenario but even then needs to be combined with weight normalisation and a top-down cognitive bias mechanism (**Appendix E**) to provide a realistic working model.

(4) Attempts have been made to use recurrent/lateral network dynamics to explain roving [34]. The systems examined in this article have all been entirely feed-forward in architecture. An intuitive understanding of the unsupervised bias hypothesis typically suggests a high-dimensional disruption of learning rather than the singular dimension shown in our analysis heretofore. This high-dimensional intuition can easily lead to mistaken interpretations of how the hypothesis may function. It is vital to remember that the bias is mathematically defined per *individual* synapse and that synapses have only a *single* dimension along which they can either grow or diminish. On the other hand, *learning* may involve improvement of performance in a dynamic high-dimensional neuro-synaptic space, and this is the source of the intuition. In fact, it is possible to imagine a recurrent dynamical system where the unsupervised bias leads to a net null learning outcome by improving performance via some synaptic weights and concurrently disimproving performance via other synapses.

This is not a very satisfactory solution however. To explain failure to learn during roving, it relies on an improbably precise balance of improving and disimproving modifications within the network. It should also be somewhat independent of the initial performance levels, since roving appears to interfere with learning across many subjects and it is improbable that all are being tested at exactly the same difficulty level with respect to their internal network organisation. Such an explanation would be more plausible if there was some suggestion of overlap in the neuronal task representations, allowing a ‘trace’ of the previous task to interfere with learning on the current one, but this is reasonably excluded via the plethora of experiments which demonstrate an absence of transfer in learning. Finally, a recurrent high-dimensional representation should typically be more, not less, likely to avoid getting trapped in a local minimum in synaptic weight space [5]. So while the unsupervised bias may theoretically lead to many non-beneficial weight changes in a recurrent network, in a high enough dimensional problem-space this bias should not impinge upon learning.

One final idea, could eventually rehabilitate the unsupervised bias hypothesis. This is the question of whether the bias can predict the statistical next choice of a subject, given their recent sequence of task performance, task presentations and task choices. The predictions of our flow field are rather strong. The question is whether it would be possible to localise the position of a human subject on the flow field (as done for simulation subjects in **Figure 9**) and then predict their performance on subsequent tasks. Since human variation in performance is quite large in bisection learning tasks it seems wholly unrealistic, at this point, that a precise positioning of the subject, in terms of performance, on the flow field can be performed.

In conclusion, upon thorough analysis, we find it hard to imagine any but the most contrived circumstances which would allow for a model built upon the unsupervised bias hypothesis to accurately model failure to learn during roving. In particular, the bias appears to have a stronger effect within a single bisection task than across multiple tasks. Everything we have explored so far indicates that, at best, there would require a complicated number of phenomena (unsupervised bias, weight normalisation, firing rate homeostasis, network architecture) which must combine in order to generate a failure to learn during roving.

In the next section we will examine other approaches to modelling of perceptual learning, and their potential applications to roving, discussing their relative merits. In **Section 4.3** we will propose our own solution to the roving problem.

### 4.2. Other models of perceptual learning

Herzog and Fahle (1997) [13] examined models of perceptual learning, for a vernier offset discrimination task, under three categories: supervised, unsupervised and reward learning models. Supervised models are largely in agreement with experimental observations, but the ability of subjects to learn even from block feedback would require an elaborate machinery of monitoring in order to deliver an appropriate supervision signal which seems unlikely (and is discounted in the paper). Furthermore, the ability of subjects to learn under conditions of only partial feedback makes the hypothesis of a supervised learning regime even more unlikely. Unsupervised models, in the definition of the paper, develop statistical internal models of input data and use this to develop classifications in the absence of any external feedback mechanism. This seems even less likely, as subjects in [13] exhibit less task improvement in the absence of feedback, and indeed disimprove in the presence of false feedback.

A hybrid model of perceptual learning, again in the vernier task, involved a feedforward network coupled to self generated feedback [14]. A set of partially tuned inputs drove the feedforward network, the feedback mechanism gated some of these inputs allowing for a sort of Hebbian type of learning with a search based selection of inputs to disable in order to maximise the difference in the outputs in a decision system. This allowed for some degree of unsupervised learning when no external feedback was provided. The system switched to using external feedback when it was available. This model still provides one of the direct implementations of a complete perceptual learning system most conceptually aligned with the idea of a biological implementation of an internal network circuit capable of selecting elements in the *actor* network for reinforcement (a supervised-unsupervised approach), without recourse to a reward learning mechanism.

One of the features of the model is that the learning rate is actively controlled by a monitoring system. Only if prior assumptions are met during the learning process, synaptic changes in the visual neurons are enabled. Otherwise, the learning rate is set to zero. For example, when more left than rightward verniers are presented, no learning occurs because the a priori assumption that both alternatives appear with the same probability is violated - in accordance with experimental data [1,15,12]. A similar situation may be at work in roving. When the unsupervised bias leads to a runaway situation, more, say, leftward responses occur and learning is disabled to avoid learning of stimuli, which are not consistent with prior assumptions.

The results presented in [13] strongly support a model based on reward learning, most likely via the modulation of a covariance-based learning rule, but this is not further explored in that paper due to the state-of-the-art limitations at that time. The main argument in favour of a reward (feedback) driven learning system is a combination of the arguments from [13], presented above, with an emphasis on biological plausibility, and indeed Occam’s razor. There are *many* specialised subsystems in the brain, so it would be foolish to argue that there is not specific feedback on a given system capable of providing the type of internal model necessary for selection of pathways for reinforcement on a particular task. But, given the highly artificial form of the stimuli typically used in perceptual learning, combined with the multiplicity of such stimuli, it seems unlikely that such high-dimensional, specific-connectivity, resources would be available for such tasks. It is simpler to assume a reward learning framework.

Despite this, it is clear that there is a multitude of learning processes at work in any one perceptual learning task. The community has worked hard to establish experimental protocols which separate these processes, but there will always be some unsupervised associative tuning, with a degree of lateral inhibition of neighbouring representations, in input layers. Top-down attention, and winner-take-all mechanisms upon choice selection, likely enhance some representations whereas others are repressed

[2], in a semi-supervised manner. A further consideration is that, learning changes the performance of the behavioural network (typically called the *actor* [28]) on a rather rapid timescale. This timescale must be matched inside the learning mechanism itself (e.g. tracking performance via a *critic* network) otherwise the system cannot maintain consistency. Since biology has largely solved the problem of perceptual learning, any model which fails in this manner does not fully explain the biology.

We have focused our work on the decision making process of the perceptual learning task. In [22] we see a dramatic early learning phase, followed by a later more gradual improvement phase. The early learning appears to be largely due to acclimatisation to the moving stimulus location. Our work can more likely be considered to model the much slower, late phase, of learning, similarly to that seen throughout in [29]. The input layer of our model can be considered to be a ‘stable’ image representation from which a decision about offset must be made. Feedback leads to learning, which leads to an improvement in the decision process, except in the case of stimulus roving. A number of other models of perceptual learning have been proposed [34, 27, 17, 30]. We will summarise them briefly here and evaluate their capacity to explain failure to learn during task roving.

Zhaoping et al. (2003) [34] developed a recurrent model of visual input processing, the testing of which led to the original work on failure to learn a bisection discrimination task during roving [22]. The model shows that a linear readout of this early processing, similar to our decision network, can lead to greater visual acuity than that directly available at the input level. This is a strong argument in favour of our architecture, justifying the level at which we place the decision circuit after image stabilisation has occurred. Ironically, despite leading to the discovery in [22] this model is not compatible with explaining roving, since no negative transfer of learning occurs in the experiments. This suggests that the roving effect is not at the input stabilisation level.

Tsodyks et al. (2004) [30] developed an influential alternative approach. They developed a cortical populations model, with Hebbian plasticity, for a contrast discrimination task. This moves the learning problem to higher cortical areas. In the words of one of the authors [31], this approach leads to difficulty when confronted with the task specificity of perceptual learning problems. Since the effects of roving are specific to certain tasks, and certain stimuli, this appears to argue against such a model.

Schäfer et al. (2009, 2007) [27, 26] developed an input tuning model, similar to that of [34], in which top-down signals [2] enhance certain parts of the receptive field. Lateral inhibition increases contrast leading to easier learning of the readout. When applied to roving, the model appears to lead to a reduced upper performance level on both tasks, when learning occurs over long timescales similar to [23]. No reduced performance is observed in the earlier learning phases. This is directly contrary to what we expect from [22, 29].

The current state-of-the-art in terms of perceptual learning modelling is perhaps the paper by Liu et al. (2015) [17]. They develop a full system based on Gabor filtered visual inputs, with varying receptive fields and orientations, which appears to explain many of the problems described in perceptual learning experiments. They have not, however, tackled failure to learn during roving. Their model is particularly interesting to us as, although it uses reward feedback to induce synaptic updates, they have encoded their update rule in a way which mathematically avoids having an unsupervised bias. In fact, their system needs only a minor modification to incorporate such a bias in the feedback signal. We have focused our model on finding the correct balance of simplicity and generalisability to answer the question as to whether the unsupervised bias hypothesis can explain roving. It would be interesting now to incorporate our ideas, which we will present in **Section 4.3**, into their model. This would presumably result in a complete model for an entire ensemble of perceptual learning experiments.

### 4.3. The single critic hypothesis

In our extended analysis we have attempted to hypothesise systems which are susceptible to the unsupervised bias hypothesis. Even in this highly contrived setting we have found it hard to construct such a system. At some point it becomes necessary to ask whether we are fighting too hard to rescue a theory, when perhaps another one could answer the problem much more simply. In fact, such a theory does exist, it just has not yet been linked to failure to learn during roving. In cognitive-behavioural analysis using fMRI it is common to test for reward prediction correlates in the brain. Recently, it has been shown, using paradigms with changing or uncertain contingencies, that the reward prediction signal appears to be normalised by the variance in the prediction [24, 25, 21].

This means that, in cases of increased uncertainty no reward prediction error (RPE) signal is produced. In a TD learning paradigm this leads to no synaptic update being performed. Such a scenario can be very easily fitted to our regime. In cases of task roving, where there is some uncertainty in the critic as to the correct reward prediction which should be produced, no reward prediction signal is created.

Furthermore, upon extreme repetition of the tasks, two separate critic representations are finally learned which immediately begin sending correct reward predictions for each of the tasks. This may also explain why in [23] it appears as though the actual learning occurs quite quickly (shortly after block 30), once it does occur.

We have implemented a simple version of this idea, with a critic which provides no feedback signal on roved tasks for the first thirty blocks. After this point, this simple critic then switches to providing task based feedback from two separate critics. A proof of concept is shown in **Figure 11**. The implementation is in keeping with models used throughout the cognitive reward learning literature [10, 9]. In practice, we might expect a slightly longer transition to learning. In future work we expect to be able to present an online critic representation learning algorithm. This will separate the critic representations based on evidence for the existence of one versus two tasks and will have a timescale of transition from a single critic to two fully separate critics.

Our hypothesis is also compatible with examples of task sequencing in perceptual learning (also referred to as roving) which did not inhibit learning [16]. This may be explained by the deliberate rhythmicity of their task switching paradigm. By embedding the task switching in a sequence the authors may, inadvertently, have provided a strong signal to the critic as to which task was being performed. This would allow separate critic representations from the first training blocks.

The single critic hypothesis is not easily experimentally distinguishable from the unsupervised bias hypothesis, both may incorporate a learning critic which leads to a sudden shift in learning. However, it has the benefit of being a much simpler theory, correlates of which have already been widely discussed in the literature.

**Figure 11:**
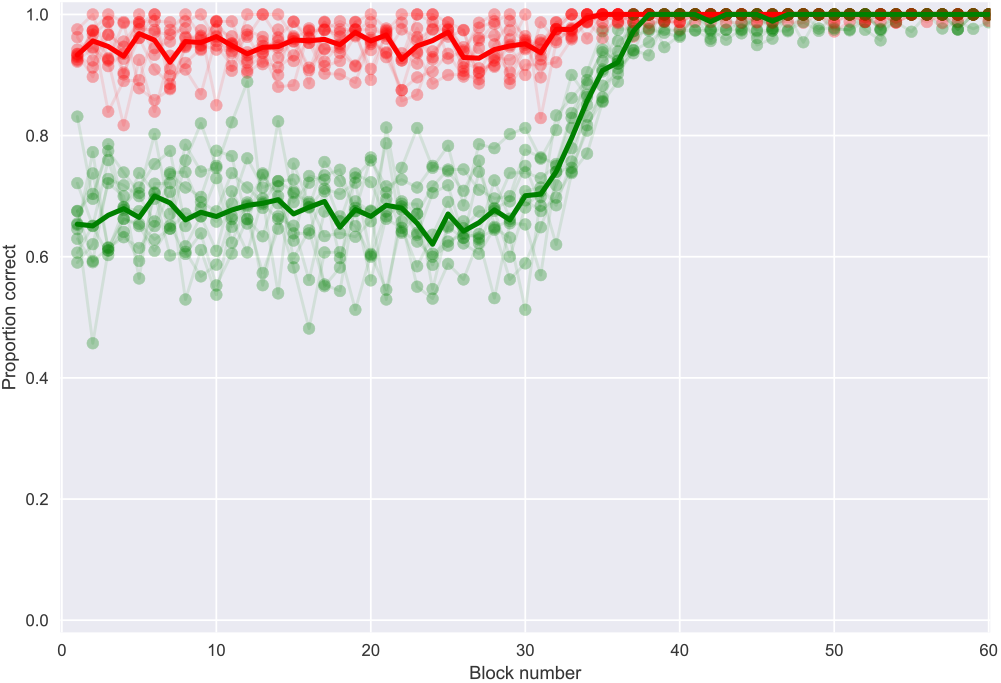
Long term task roving eventually leads to learning under the single critic hypothesis. For the initial 30 blocks no weight updates are performed. From training block 31 two separate critics are used, one for the ‘**easy**’ task (red) and the other for the ‘**hard**’ task (green). Performance in both tasks quickly improves until saturation. Note, we needed to use a simple mechanism of top-down 50:50 left-right expectation to overcome the internal unsupervised biases which would otherwise have prevented learning. This mechanism was enabled throughout all blocks.

## 5. Conclusion

Any good theory for explaining failure to learn during bisection task stimulus roving must fulfill a number of requirements. The model must learn, consistently, upon block presentation of a single left-right bisection task. It must then, fail to learn upon a comparable number of presentations of two tasks randomly alternating (roved) on a trial-to-trial basis. It must perform this failure to learn, while still responding effectively 50% of the time with either outcome, similarly to human subjects. Finally, a full theory of roving should have a mechanism which explains the ultimate success in learning, after at least one order of magnitude more trials, as shown in [23].

In this paper, we have presented the unsupervised bias hypothesis for explaining the failure to learn during task roving [11]. We then presented a general mathematical modelling approach suitable to the analysis of the original hypothesis, before extending the analysis to explain how the models originally presented actually functioned. Finally, we have analysed the unsupervised bias hypothesis in its entirety, as to whether any alternative implementations of the architecture could be conceived as supporting the hypothesis. Our approach has combined aspects of logic, analytical mathematics, numerical mathematics and finally simulations.

Ultimately, we find that any simple or intuitive understanding of the unsupervised bias hypothesis fails when it comes to explaining failure to learn during roving. The mechanism by which such simplistic intuitions typically fail is the fact that synapses can only increase or decrease their weights in a single dimension. The hypothesis makes a statement of the effects of the error in reward prediction at the level of single synapses. Due to the single dimension in which the synapse operates, this can only lead to over-shoot or shortfall in the update. This is particularly relevant in a case such as perceptual learning where all presentation ratios are equal and constant.

The failure of the unsupervised bias hypothesis to explain failure to learn during task roving should not be construed as a rejection of the existence of unsupervised biases throughout the brain [18, 8]. This is an important topic. However, the results in this paper indicate that such biases are likely to either slow learning or lead to an inability to settle on the perfect solution, rather than actively working against learning. This is particularly consistent with the work presented in [18]. Here, we add to that understanding most importantly by our inclusion of neuronal task representation in the analysis. Indeed, what was particularly interesting in examining this hypothesis is how the mechanisms of the brain naturally counteract such potential biases. Furthermore, in a high-dimensional space these effects are likely to be negligible [5] and are more likely artefacts of the low-dimensional limitations of the simulation studies to date.

Potentially a far more interesting aspect of the unsupervised bias is its ability to perform a natural symmetry breaking operation in the brain. The tendency to have difficulties, in our scheme, in encoding both ‘left’ and ‘right’ should lead over time, even in the presence of mechanisms such as weight normalisation, etc., to specialisation towards one stimulus or the other. This could be yet another natural mechanism towards the development of input specific neuronal tuning curves over time.

In this paper we have focused on the existing primary hypothesis for explaining roving. To do so, we have developed a model which incorporates all of the mechanisms mentioned in this paper and more. What remains to be done is to develop a full generic theory of critic representation learning. This can then provide reward predictions normalised by the uncertainty in the prediction to the model. Work is currently underway on this task. This will not, however, modify any of the existing experimental predictions of perceptual task roving. What is of greater interest now is to see whether an fMRI experiment, involving roving, can pick up on this variance normalised critic, but this must be tested in the case of exceedingly long numbers of trials as done in [23].

Theory has long posited the idea of a critic in reward learning. Experimentally separating the task performance network (the actor) from the reward predictor network (the critic) has rarely been possible. Roving may be one of the first, human subject experimental paradigms, which could allow us to examine features of the critic, such as: what the metric of task similarity is for the critic; and how task similarity modifies the critic learning timescale. The realisation that a single critic is responsible for the failure to learn during roving is a vital step in this process.

## Appendix A. Mathematics of analysis of simple network

Continued from **Section 3.1** we can also analyse the dynamics of the synapses on the ‘**hard**’ task.

We will now proceed to the ‘**hard**’ task example, to see if this is also learnable under the weight update scheme. Upon presentation of a negative input we again expect Δ*w*_12_ < 0 for a correct ‘Left’ choice, and Δ*w*_22_ > 0 for correct classification on a positive input, ‘Right’ choice. Indeed, close examination of Table 2 will show that the only differences in the update rules for the ‘hard’ task are in the *Δ* and *R^true^* values used.

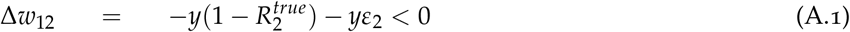

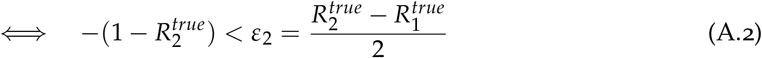

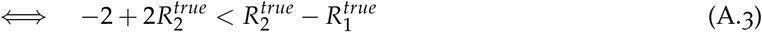

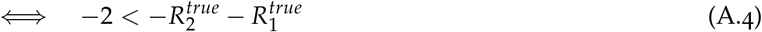

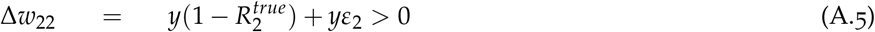

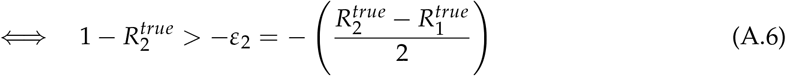

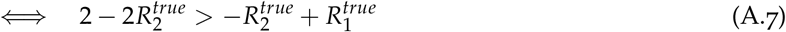

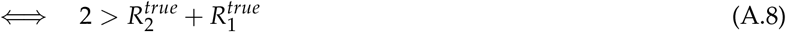

And for the incorrect classifications (error trials), we still want the weights to move in the same directions,

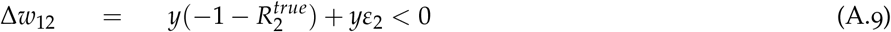

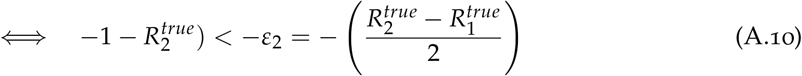

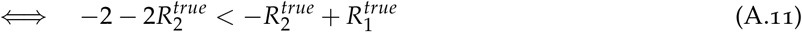

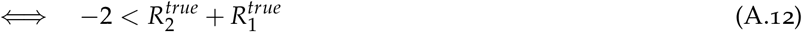

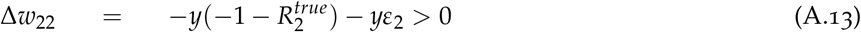

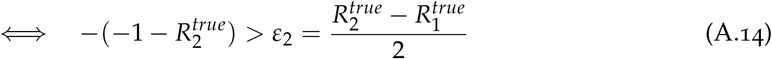

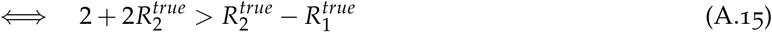

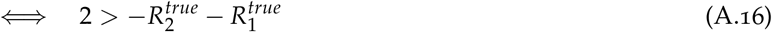

In each of the cases above, 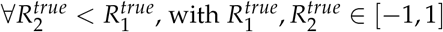, the learning conditions hold. This means that in each and every case, learning *works* and is not overpowered by the unsupervised bias.

## Appendix B. Mathematics of Linear Output Encoding

In the simple model of perceptual learning presented in **Sections 2.1** and **3.2** the system chooses ‘Left’ or ‘Right’ based on which output neuron has the higher firing rate. Here we will develop a mathematical treatment, which will allow us to analyse the trajectories in probability space of correctly identifying a ‘Left’ or ‘Right’ stimulus on a given task. These trajectories will allow us to separately distinguish the effects of the **pure learning, internal bias** and **external bias** terms, presented in **Section 3.2.2**. For this analysis we will relax the winner-takes-all (WTA) requirement, on the output encoding, of the Herzog et al. [11] model. Such a mechanism can, technically, be treated analytically but it requires us to fix certain assumptions about firing rates which increase the complexity of the analysis, without bringing greater insight.

Choosing ‘Left’ or ‘Right’ for a given stimulus is a result of a combination of the input representation, the network weights and the trial noise. We can define,

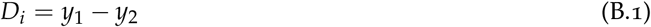

as the difference in the output firing rates, in response to stimulus *i*. Note, all values should be parameterised by time, t, which we will omit here for readability. The value of *D_i_* is probabilistic and we can write the probability of choosing output 1 (‘Left’) for input *i* as,

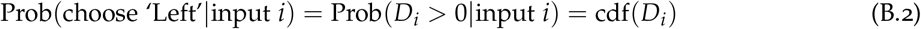

where cdf() is the cumulative density function. For a basic understanding, we note in passing that cdf(0.0) = 0.5 corresponds to the expectation of both outputs being essentially indistinguishable and noise will drive the choice on an individual trial with equal likelihood. We can further transform *D_i_* into *D+* which we will define as,

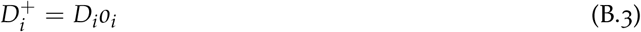

where *o_i_* is the correlation between *y*_1_ > y_2_ and *R*_i_. What this means is that D+ is always positively correlated with reward and takes account for the fact that output 2 should be active for ‘Right’ stimuli. We will define,

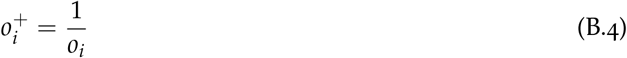

giving us

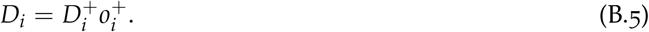

If we write the weight change rule as follows,

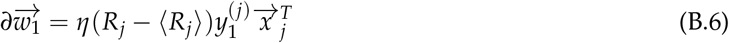

where the subscript 1 denotes all weights targeting output neuron 1 and *j* is the input identity. Then we can write the trial averaged weight change as,

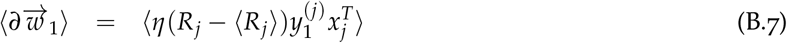

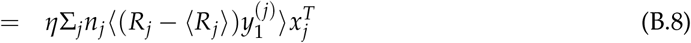

Similarly, we can write out the dynamics of *D*,

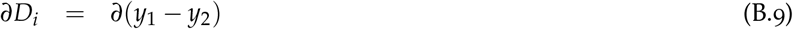

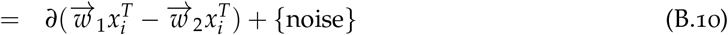

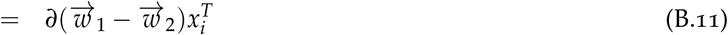

Now, we can look at the trial and noise averaged change in *D* via,

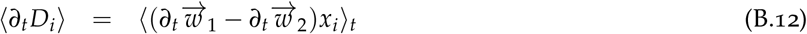

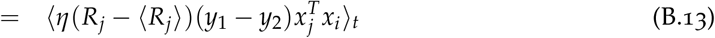

This allows us to define a *similarity matrix*

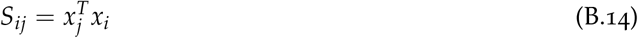

which represents the task representation similarity, in the input layer, between tasks *i* and j. We can then continue,

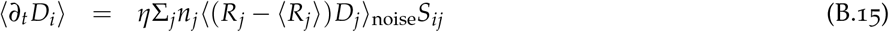

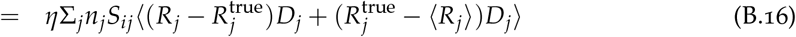

where 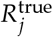 is the true expectation of reward on task *j*, which may differ from the running average expectation 〈*R_j_*〉. Then,

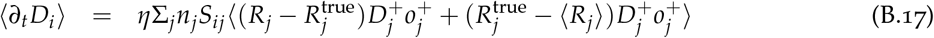

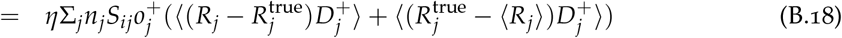

We can define,

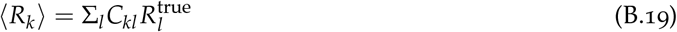

then if *A* = *I* – *C*, where *I* is the identity matrix, we obtain,

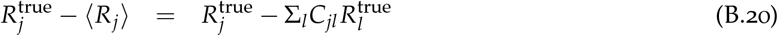

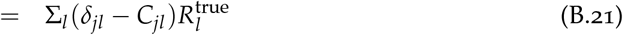

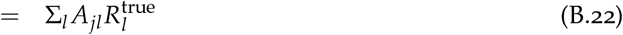

We call *C* a *confusion matrix* as it defines the proportions to which each task contributes to the reward prediction critic. We assume that the critic maintains perfect approximations to 〈*R*〉, made up of the component true averages 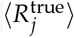.

Combining this formulation, with a result from [32] for the differentiation we obtain,

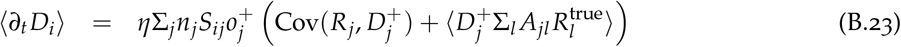

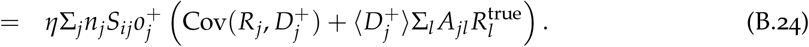

In our system, 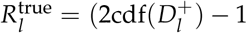 by construction. And, the result from [32] gives us,

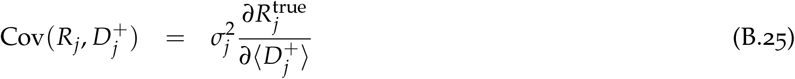

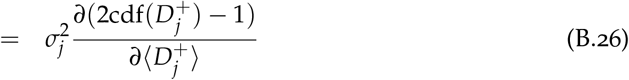

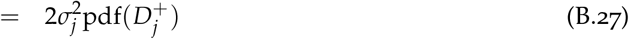

The above results in an equation which we can use for our flow field plots,

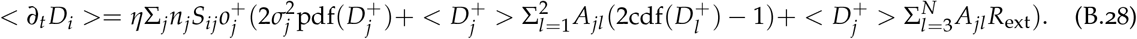

Here we have separated the first two contributors to the critic from the subsequent *N* — 2 contributors. This corresponds to our task setup, where only the first two contributors share in the task representation for task i. The first term, containing 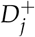 corresponds to the pure learning; the second term, 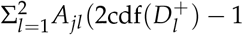, to the **internal bias**; the final term, 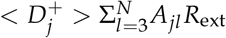, to the **external bias**.

For a probabilistic plotting, as used in the figures, we observe that,

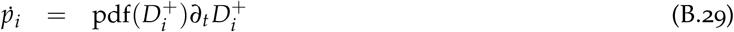

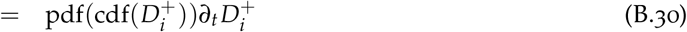

## Appendix C. Mathematics of Binary Output Encoding

This follows the same form of derivation as for Linear outputs, but restricts the firing rates to binary 1 or 0. Only one output is allowed to be active upon classification of a given task. For binary output neurons, we can derive the system performance dynamics by recognising that

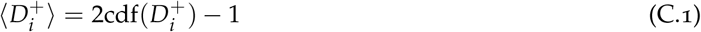

and following a similar set of operations as for the linear case, we obtain,

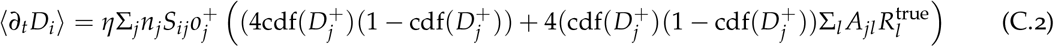

It is also possible to combine the mathematics from Appendix B and Appendix C to produce a mathematics of a winner-takes-all linear output encoding, but it requires an assumption of what the mean output firing rate will be across all probabilities of correctly classifying the task. This method is also applicable to the case of weight or firing rate normalisation. It is however, beyond the useful scope of this paper.

## Appendix D. Expanding the Output Population Encoding

In order to expand the output population encoding, while maintaining the synaptic nature of the weight update rule, we have developed a simple mathematical expression which gives separate single neuron and population firing rates on a given task. This means that whereas classification decisions are made by comparing the population firing rates (two populations, one ‘Left’ and one ‘Right’), the weight update rule remains based on the single post synaptic neuronal firing rates.

For every trial, we have two noise terms,

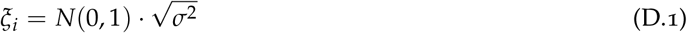

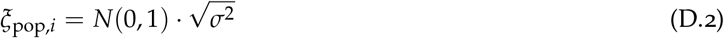

If we are working with a, per task, output population encoding size *N* > 1 then we calculate two firing rates,

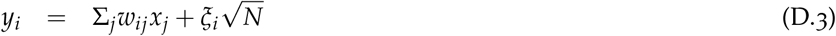

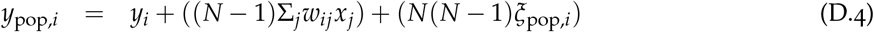

Winner-takes-all is applied, where appropriate, after the population firing rate has been calculateD This formulation allows us to modify the output population size while maintaining a constant signal-to-noise ratio.

## Appendix E. Simulation model description

We take the model presented in [11] as our base network, then add a number of optional features which allow us to better probe different aspects of the unsupervised bias hypothesis for failure to learn during task roving. Much of this is described in the text, but we will provide a summary here.

The network is a two-layer feedforward rate-based neuronal network. Inputs are separated by task, with 50 input neurons for each task. Input representation is linear, based on distance from the true bisector, *φ*, *x_j_*(*φ*) = *a_j_* + *b_j_*.*φ*. *a* ~ *N*(2,0.5), *b* ~ *N*(0, *β*), where *β* is 0.25 and 0.375 for tasks 1 and 2 respectively. The larger value of *β* corresponds to the ‘Easy’ task in the text. Initial weight values are setup according to *W_ij_* = *U*(0,1) + 2*b_i_*(− 1)^j^. Postsynaptic firing rates are described by, *y_i_* = Σ*_i_W_ij_x_j_* + *ξ_i_* unless the expanded output population encoding is used, in which case Appendix D describes the equations. A winner-takes-all mechanism is typically appied to the output firing rates, which sets the firing rate of the non-chosen option to 0.

Reward, on trial *n*, *R*^(*n*)^ is given as a result of correct, *R*^(*n*)^ = 1, and incorrect, *R*^(*n*)^ = −1, input classification. A running average of reward, 〈*R*〉 is maintained,

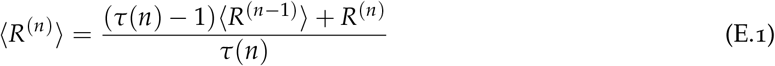

where *τ*(*n*) = min(50, *n*). Weight updates are performed, following application of winner-takes-all, after each trial via,

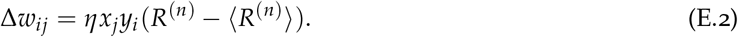

For comparisons with [11] we typically include 80 trials in a block and 14 blocks in an experiment. The learning rate *η* = 0.002 except for comparisons with deterministic limit of mathematics, in which case it was reduced to *η* = 0.0005.

**Figure E 12:**
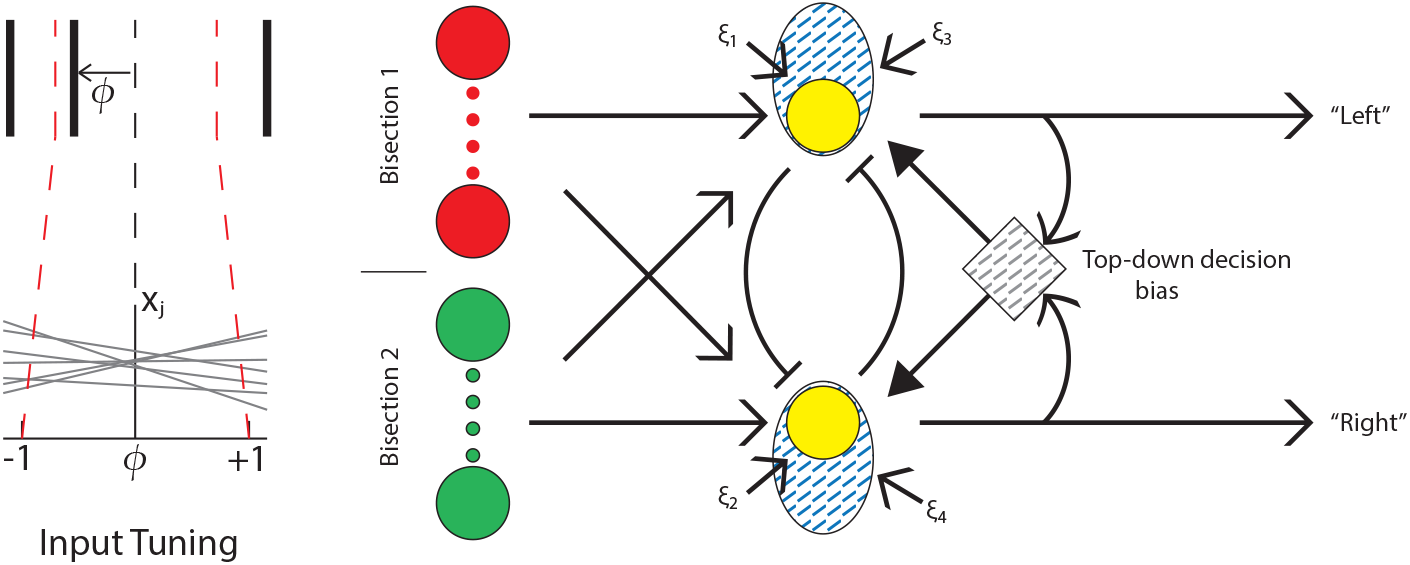
Our full network implementation incorporates the possibility of an expanded output population encoding and also top-down feedback which drives a 50:50 left-right bias (*we have experiments to show this bias*). Not shown in the diagram are the multiple (optional) mechanisms for weight and output firing rate normalisation.

In addition to the main analysis provided in the Results, we also explored the effects of weight normalisation and a top-down cognitive bias, combined with an unsupervised bias, on task learning. We implemented both multiplicative and subtractive normalisation, largely focusing on multiplicative normalisation via,

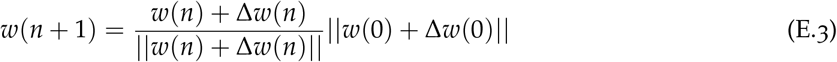

We maintain the scaling of the initial weight distribution, via *w*(0) for stability. We further rescale the norm by the number of neurons for direct comparison across network sizes, such that,

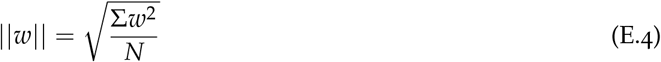

where *N* is the number of weights (synapses).

We explored a number of mechanisms for top-down bias in order to maintain the human expected choice ratio of 50:50 left - right. The principal mechanism, is a simple choice integrator which differentially subtracts/adds from the output neuron which has been over-represented/under-represented in the recent history of task choices.

## Acknowledgements

DH wants to give special thanks to Guillermo Aguilar and Alessandro Barri for their feedback on early versions of this document. Henning Sprekeler was involved in the development of the mathematical treatment in this paper, but has declined the opportunity to be co-author.

## Bibliography

[1] K. C. Aberg and M. H. Herzog. Different types of feedback change decision criterion and sensitivity differently in perceptual learning. Journal of Vision, 12(3):3–3, 2012.

[2] Merav Ahissar and Shaul Hochstein. The reverse hierarchy theory of visual perceptual learning. Trends in Cognitive Sciences, 8(10)457–464, October 2004.

[3] Karlene Ball and Robert Sekuler. Direction-specific improvement in motion discrimination. Vision Research, 27(6)953–965, 1987.

[4] Roy E. Crist, Mitesh K. Kapadia, Gerald Westheimer, and Charles D. Gilbert. Perceptual learning of spatial localization: Specificity for orientation, position, and context. Journal of Neurophysiology, 78(6)2889–2894, 1997.

[5] Yann N. Dauphin, Razvan Pascanu, Caglar Gulcehre, Kyunghyun Cho, Surya Ganguli, and Yoshua Bengio. Identifying and attacking the saddle point problem in high-dimensional non-convex optimization. In Advances in neural information processing systems, pages 2933–2941, 2014.

[6] Barbara Anne Dosher and Zhong-Lin Lu. Perceptual learning reflects external noise filtering and internal noise reduction through channel reweighting. PNAS, 95(23):13988–13993, 1998.

[7] Manfred Fahle. Perceptual learning: specificity versus generalization. Current opinion in neurobiology, 15(2):154–160, 2005.

[8] Nicolas Frémaux, Henning Sprekeler, and Wulfram Gerstner. Functional requirements for reward-modulated spike-timing-dependent plasticity. The Journal of Neuroscience, 30(40):13326–13337, June 2010.

[9] Samuel J. Gershman, Marie-H. Monfils, Kenneth A. Norman, and Yael Niv. The computational nature of memory modification. eLife, 6:e23763, 2017.

[10] Samuel J. Gershman and Yael Niv. Learning latent structure: carving nature at its joints. Current opinion in neurobiology, 20(2):251–256, 2010.

[11] Michael H. Herzog, Kristoffer C. Aberg, Nicolas Frémaux, Wulfram Gerstner, and Henning Sprekeler. Perceptual learning, roving and the unsupervised bias. Vision Research, 61:95–99, May 2012.

[12] Michael H. Herzog, Knut RF Ewald, Frouke Hermens, and Manfred Fahle. Reverse feedback induces position and orientation specific changes. Vision research, 46(22):3761–3770, 2006.

[13] Michael H. Herzog and Manfred Fahle. The role of feedback in learning a vernier discrimination task. Vision Research, 37(15):2133–2141, August 1997.

[14] Michael H. Herzog and Manfred Fahle. Modeling perceptual learning: difficulties and how they can be overcome. Biological Cybernetics, 78(2):107–117, February 1998.

[15] Michael H Herzog and Manfred Fahle. Effects of biased feedback on learning and deciding in a vernier discrimination task. Vision Research, 39(25):4232–4243, 1999.

[16] Shu-Guang Kuai, Jun-Yun Zhang, Stanley A. Klein, Dennis M. Levi, and Cong Yu. The essential role of stimulus temporal patterning in enabling perceptual learning. Nature Neuroscience, 8(11):1497–1499, 2005.

[17] Jiajuan Liu, Barbara Anne Dosher, and Zhong-Lin Lu. Augmented hebbian reweighting accounts for accuracy and induced bias in perceptual learning with reverse feedback. Journal of vision, 15(10):10–10, 2015.

[18] Yonatan Loewenstein. Robustness of learning that is based on covariance-driven synaptic plasticity. PLoS Comput Biol, 4(3):e1000007, 2008.

[19] Z.-L. Lu and B. A. Dosher. Perceptual learning retunes the perceptual template in foveal orientation identification. Journal of Vision, 4(1):5–5, 2004.

[20] Suzanne P. McKee and Gerald Westhe. Improvement in vernier acuity with practice. Attention, Perception, & Psychophysics, 24(3):258–262, 1978.

[21] Yael Niv, Jeffrey A. Edlund, Peter Dayan, and John P. O’Doherty. Neural prediction errors reveal a risk-sensitive reinforcement-learning process in the human brain. Journal of Neuroscience, 32(2):551–562, 2012.

[22] Thomas U. Otto, Michael H. Herzog, Manfred Fahle, and Li Zhaoping. Perceptual learning with spatial uncertainties. Vision Research, 46(19):3223–3233, October 2006.

[23] Khatuna Parkosadze, Thomas U. Otto, Maka Malania, Archil Kezeli, and Michael H. Herzog. Perceptual learning of bisection stimuli under roving: Slow and largely specific. Journal of Vision, 8(1):5, November 2008.

[24] Kerstin Preuschoff, Peter Bossaerts, and Steven R. Quartz. Neural differentiation of expected reward and risk in human subcortical structures. Neuron, 51(3):381–390, 2006.

[25] Kerstin Preuschoff, Steven R. Quartz, and Peter Bossaerts. Human insula activation reflects risk prediction errors as well as risk. Journal of Neuroscience, 28(11):2745–2752, 2008.

[26] Roland Schäfer, Eleni Vasilaki, and Walter Senn. Perceptual learning via modification of cortical top-down signals. PLoS Computational Biology, 3(8):e165, 2007.

[27] Roland Schäfer, Eleni Vasilaki, and Walter Senn. Adaptive gain modulation in v1 explains contextual modifications during bisection learning. PLOS Comput Biol, 5(12):e1000617, December 2009.

[28] R.S. Sutton and A.G. Barto. Reinforcement learning: An introduction. IEEE Transactions on Neural Networks, 9(5):1054–1054, September 1998.

[29] Elisa M. Tartaglia, Kristoffer C. Aberg, and Michael H. Herzog. Perceptual learning and roving: Stimulus types and overlapping neural populations. Vision Research, 49(11):1420–1427, June 2009.

[30] Misha Tsodyks, Yael Adini, and Dov Sagi. Associative learning in early vision. Neural Networks, 17(5âĂŞ6):823–832, June 2004.

[31] Misha Tsodyks and Charles Gilbert. Neural networks and perceptual learning. Nature, 431(7010):775–781, 2004.

[32] Ronald J. Williams. Simple statistical gradient-following algorithms for connectionist reinforcement learning. Machine Learning, 8(3):229–256, 1992.

[33] Cong Yu, Stanley A. Klein, and Dennis M. Levi. Perceptual learning in contrast discrimination and the (minimal) role of context. Journal of Vision, 4(3):4, 2004.

[34] L. Zhaoping, Michael H. Herzog, and Peter Dayan. Nonlinear ideal observation and recurrent preprocessing in perceptual learning. Network: Computation in Neural Systems, 14(2):233–247, January 2003.

